# Invariant synaptic density across species

**DOI:** 10.1101/2024.07.18.604056

**Authors:** André Ferreira Castro, Albert Cardona

## Abstract

The nervous system scales with animal size while preserving function, yet the principles underlying this stability remain unclear. Here, we analyse the ultra-structure and connectivity of thousands of neuronal cells across species, including fly, zebrafish, mouse, and human, and found a conserved feature that stabilises neuronal responses across scales: an average of one synapse per micrometre of dendritic cable. We show that the appropriate synaptic density is shaped by correct axon-dendrite positioning and synaptic transmission during development. We find that this specific synaptic density is linked to wiring optimisation in neurons, where dendrites minimise cable length and conduction delays. Finally, simulations indicate invariant synaptic density as a neuronal design principle, conserved for its ability to synergise with other cell-intrinsic properties to stabilise voltage responses across cell types and species.

## 1 Introduction

Body size varies by 21 orders of magnitude in the animal kingdom [1]. In contrast to most animal organs, which scale through the proliferation of dividing cells of similar size, brains expand by modulating the size of non-dividing neuronal cells to innervate space during development [2, 3]. As a result, there is notable diversity in neuronal size and structure within individual brains and between different species [4–6]. Given that size inherently influences the rates of all biological processes and structures, from cellular metabolism to information processing [7, 8], a crucial and unresolved question in neuroscience is how neurons maintain essential computational functions while optimising resources, despite size variations [9–11].

In various domains of biology and ecology, constraints on organism function and design are often understood through allometric scaling relationships, which reveal predictable patterns in how biological traits change with size [12, 13]. Within this framework, invariant properties, features that remain constant between species, underlie functions that must be preserved and emerge as fundamental constraints that shape and refine these scaling patterns [1, 9]. Through this lens, biological traits adapt within the constraints of scaling and invariance, revealing universal patterns in the organisation of life while providing normative explanations for these patterns across scales [14, 15].

In neuroscience, morphological scaling laws have long been used to describe the structure of dendritic arbours [16–18]. One prominent example is a 2/3 power-law relationship, which predicts how dendritic length, spanning volume, and synapse number scale relative to one another under principles of wiring optimisation, such as minimisation of cable length and conduction delay [19]. Remarkably, even neurons from the same cell-type can differ in size by several orders of magnitude across developmental stages and species [19–22]. From cable theory alone, such variation in dendritic size is expected to alter membrane properties, including input resistance and voltage attenuation, which affect synaptic integration and overall cell excitability [23–25]. Nonetheless, conserved neuronal types often exhibit strikingly similar firing rates and voltage responses [26]. This apparent mismatch between morphological divergence and functional stability poses a fundamental problem: how do neurons maintain consistent synaptic integration despite large variations in size and cable properties [27, 28]?

To address this question, several scaling principles have been proposed to explain how neurons preserve electrotonic properties despite morphological variation [27, 28]. More recently, a general principle, dendritic constancy, has been proposed, suggesting that neurons can preserve stable voltage responses and firing rates by maintaining a constant synaptic density along dendrites, despite variations in morphology [29]. However, this principle appears in conflict with evidence that synapse density varies across cell types and brain areas [30–32]. Establishing whether dendritic synaptic density conservation represents a general biological constraint or a context-dependent adaptation is critical for understanding how dendritic structure maintains stable computation across neuronal systems.

Testing scaling principles experimentally remains challenging due to limited ultra-structural data on dendritic trees and their connectivity. Although neuroscience has recently entered a “post-connectome” era, with detailed connectomes now available for species ranging from nematodes to humans [33–37], few datasets include fully reconstructed dendritic trees with all synapses attached. Reconstructing these connectomes requires extensive proofreading, and current AI pipelines often fall short, yielding variable completeness rates in dendritic and synapse attachment across circuits and species [38–40]. For example, dendrite completeness rates vary widely: from approximately 75–100% in the *Drosophila* larva brain connectome [33], to 20–90% in the Hemibrain project [41], 47% in FlyWire [34], and around 67% in the human cortex [37]. Although these connectomes enable detailed species-specific insights, such as predicting visual functions in *Drosophila* based on anatomy alone [42]—their capacity to reveal universal principles of neuronal organisation remains limited [43]. As a result, comparisons of neuronal scaling and function across species are still in early stages, leaving fundamental questions about the structure-function relationship unanswered [44].

Here, to address this gap, we processed and analysed thousands of densely reconstructed single-cell morphologies and synaptic structures across EM-derived connectomes from *Drosophila*, zebrafish, mouse, and human to explore how neuronal structure scales with function. In line with the dendritic constancy principle, we found that dendritic cable synapse density (hereafter synaptic density) remains remarkably invariant, converging to one synapse per micrometre across species despite varying by 5 orders of magnitude in cable length. Analysis of genetically modified *Drosophila* shows partner location and synaptic transmission shape synaptic density during development. We also confirm the quantitative match of the real morphological data and the 2/3 power-law wiring optimisation scaling relationship. To directly link synaptic density invariance and wiring optimisation scaling, we performed a sensitivity analysis using this law, providing evidence that density invariance may arise from structural pressures to minimise wiring cost. Finally, we show through simulations that synaptic density invariance, in synergy with dendritic radius and synaptic size, can lead to stable voltage responses across a wide range of dendritic trees and species. Our findings suggest that the convergence in synaptic density serves as a functional backbone for computational stability, while linking principles of resource optimisation and functional stability across evolutionary and developmental scales.

## 2 Results

### 2.1 Synaptic density converges to 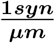 across species

To investigate synaptic density distributions across species, we processed, analysed and annotated thousands of single-cell morphologies from multiple synaptic-level resolution electronic microscopy datasets [33, 41, 45–47]. As shown in **Figure 1A-B**, the number of synapses across different cell types in *Drosophila*, zebrafish, and mouse nervous systems scales nearly linearly with dendritic length. The log–log regression shows a strong fit (*r*^2^ = 0.96; *slope* = 1.07), and close agreement with the unity line (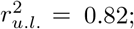 ; *slope* = 1), consistent with an average synaptic density of approximately 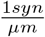, despite large differences in cell size.

**Fig. 1.**
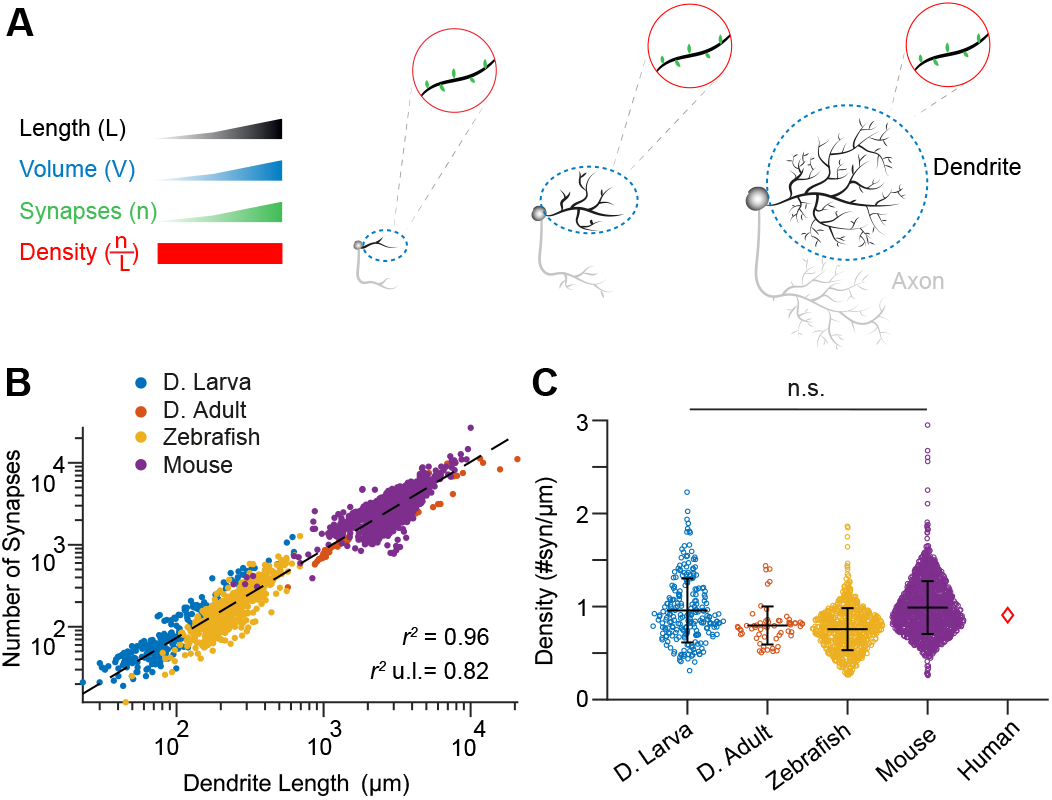
Synaptic density comparisons across species. **A**, Schematic of the quantities used throughout this study. Namely, dendritic length (*L*; black), volume (*V* ; dashed blue ellipse), and number of synapses (*n*; green). Inset zooming in dendrite cable in red highlights that the dendritic synaptic density is constant regardless of cell size or complexity. Axon is depicted in gray and soma as the circular gray structure connecting dendrites and axons. **B**, Log–log plot showing the relationship between total dendritic length and the number of synapses per neuron across four species: *Drosophila* larva (blue), adult *Drosophila* (orange), zebrafish (yellow), and mouse (purple). Each dot represents a single reconstructed neuron (*N*_*Total*_ = 2486, *N*_*D*.*Larva*_ = 233, *N*_*D*.*Adult*_ = 60, *N*_*Zebrafish*_ = 709, *N*_*Mouse*_ = 1484; see Table 1, 2 and Methods 3.5). A linear fit (dashed line) across all data reveals a strong scaling relationship (*r*^2^ = 0.96; *slope* = 1.07), closely matching the unity line (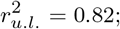; *slope* = 1). **C**, Swarm plot of synaptic densities across species. Each data point is one neuron, with the colour corresponding to species. Horizontal black line indicates mean values per species; error bars, standard deviation. Red diamond shape indicates mean values of corresponding species from [44]. Pair-wise statistical analysis using permutation tests (10000 permutations), “n.s.” non significant.

To further examine species-specific patterns in synaptic density, we next analysed the distribution of individual neuron values within each dataset (**Figure 1C**). Interestingly, this convergence persists despite variations in dendritic lengths, circuits, and ecological niches (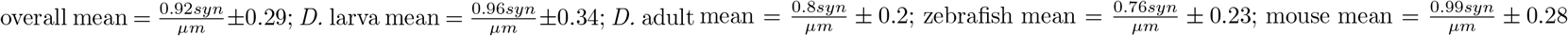; **Table 1**; **Figure S1**; see Methods 3.5 for the specific lower density in zebrafish, which in all likelihood emerges from an underdetection of the number of synapses). We also show in SI that synapses appear homogeneously distributed along dendrites in across all datasets (see Methods 3.5; **Table 2**). In particular, in the most densely reconstructed datasets, specifically *Drosophila* larva and mouse, the correspondence with 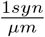 is remarkably robust, with no statistical difference found between them (difference of the 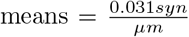, permutation test *p* = 0.13). The comparison between *Drosophila* larva and mouse was chosen because these datasets offer the most complete reconstructions, enabling robust statistical analysis. Adding human data from a recent study further substantiates this pattern (data from [44], mean 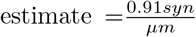; **Figure 1C**). The convergence in synaptic density is unexpected considering that earlier volumetric studies, which estimate synapse density per unit volume without direct ultrastructural reconstruction, revealed significant variations in synapse density across circuits and species [30–32]. Taken together, these results suggest that, within the framework of the dendritic constancy principle, convergence in synaptic density may act as a functional backbone to support stable voltage responses to synaptic input across scales.

**Table 1.**
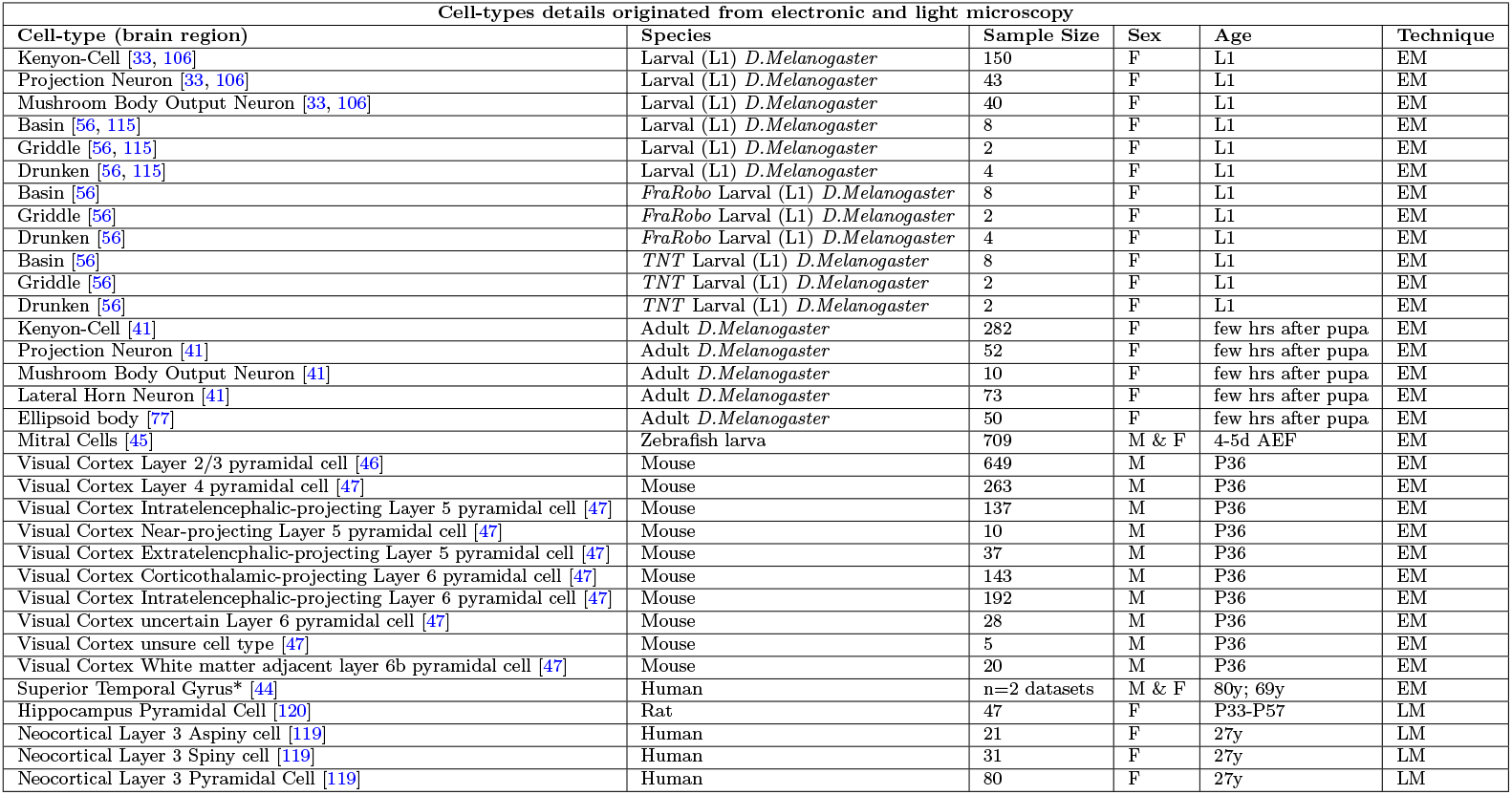
Details of species, brain regions and cell-types used in this study originated from electronic (EM) and light (LM) microscopy. The table provides a comprehensive overview of the cell-types investigated across various species, including *Drosophila* (both larval and adult stages), zebrafish, mouse, rat and human. For each cell-type, the corresponding species, sample size, sex, and age are listed. Sex is Female (F), Male (M). Age is hours after egg laying (AEL), hours after pupating, post embryonic days (*P*_*days*_), or human subjects age years (y). Star sign denotes datasets from which estimations of synaptic density were retrieved, but anatomical data were not available. References for the data sources are provided.

**Table 2.** Quantitative summary of synaptic properties across different species. This table provides a quantitative summary of synaptic properties across various species, detailing synaptic density, inter-synaptic distance, and spatial location. The values are presented as mean ± standard deviation (SD). Spatial location, indicating variability in synapse positioning. The location ranges between [0 − 1], with value 0 meaning the most proximal, 0.5 is uniformly distributed at random, and 1 the most distal (see Methods 3.5). Star sign denotes corrected values (uncorrected zebrafish synaptic 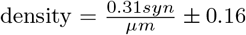; uncorrected zebrafish inter-synaptic 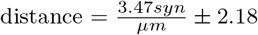; see Methods 3.5)

| Synaptic quantifications details |  |  |  |
| --- | --- | --- | --- |
| Species | Synapse density<br>(mean ± SD) | Inter-synaptic distance<br>(mean ± SD) | Spatial location<br>(mean ± SD) |
| *Drosophila* larva | $\frac{0.96syn}{\mu m} \pm 0.34$ | $\frac{1.31syn}{\mu m} \pm 0.49$ | $0.54 \pm 0.06$ |
| *Drosophila* adult | $\frac{0.81syn}{\mu m} \pm 0.22$ | $\frac{1.89syn}{\mu m} \pm 0.51$ | $0.66 \pm 0.16$ |
| Zebrafish adult | $\frac{0.76syn*}{\mu m} \pm 0.23$ | $\frac{1.23syn*}{\mu m} \pm 0.47$ | $0.31 \pm 0.1476$ |
| Mouse adult | $\frac{0.99syn}{\mu m} \pm 0.28$ | $\frac{1.57syn}{\mu m} \pm 0.45$ | $0.45 \pm 0.07$ |

### 2.2 Partner position and synaptic transmission shape synaptic density during development

The observed convergence of synaptic density to approximately 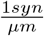 across species raises the question of how this feature is established during neuronal development. Is it a passive consequence of cell growth, or does it require active regulatory processes? Prior work in *Drosophila* [48–50], *C. elegans* [51] and mouse [52] have suggested that synaptic density during development is conserved, despite steep increases in the number of synapses throughout development. However, the underlying mechanisms responsible for this conservation remain uncharacterised.

Synaptogenesis is thought to depend on molecular recognition cues, dendritic morphology, and activity of presynaptic partners, all of which are likely to influence how synaptic density is established and maintained during development [53–56]. To investigate the contribution of these factors, we analysed neuronal morphologies of wild-type and genetically perturbed mechanosensory neurons within the *Drosophila* larval ventral nerve cord (VNC), reconstructed from EM data [56]. We selected this dataset because it is, to our knowledge, the only available EM volumes that include both wild-type and mutant neurons relevant to our question within the same identified circuit. These mutants carry genetic constructs that either shift arbour locations (*Mechano-FraRobo*) or suppress presynaptic exocytosis (*Mechano-TNT* ; **Figure 2A**; see Methods 3.5). We quantified synaptic density in these neurons to assess how altered partner access or synaptic transmission impacts synapse allocation.

**Fig. 2.**
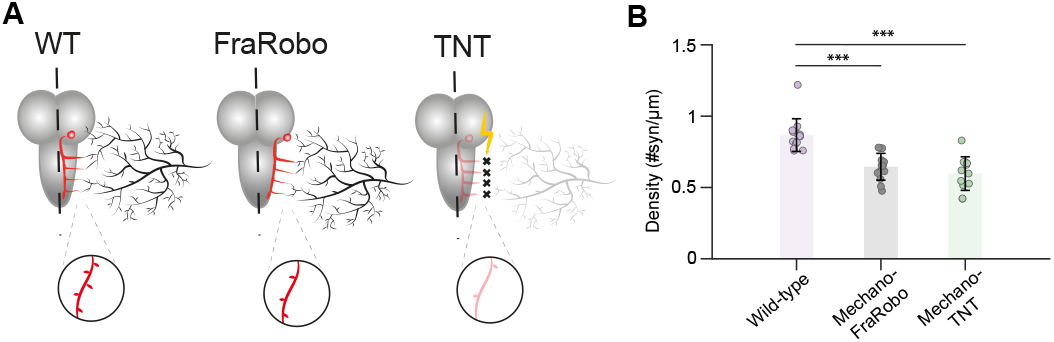
Synaptic density comparisons across *Drosophila* mutants. **A**, Schematic of EM volumes used in the comparative dendritic synaptic density analysis. Left scheme, wild-type: depicts the larval *Drosophila* brain and ventral nerve cord (grey) with a central dashed line indicating the body’s axis of left-right symmetry. The mechanosensory axon is shown in red, connecting to dendritic trees (black). The inset highlights synaptic density. Central scheme, FraRobo mutant: similar to WT scheme, but with the mechanosensory axon (red) shifted from its typical location. Right scheme, TNT mutant: illustrates synaptic transmission supression in the mechanosensory axons **B**, Bar plot of synaptic densities across wild-type *D*. larvae mechanosensory neurons and mutants. On each plot, coloured bars corresponds to mean values, respectively per EM volume. Error bars indicate standard deviation (*N*_*WT*_ = 14, *N*_*TNT*_ = 12, *N*_*FraRobo*_ = 14; see Methods 3.5). Pair-wise statistical analysis using permutation tests (10000 permutations), ∗ ∗ ∗ indicates *p <* 0.001.

We found that while wild-type morphologies exhibit synaptic densities close to 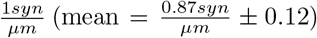, both mutants have significantly lower densities (*Mechano-FraRobo* 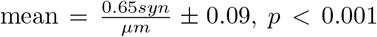; *Mechano-TNT* 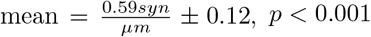; **Figure 2B**), indicating that disruptions in partner positioning or synaptic transmission impacts synaptic formation. Importantly, despite large and significant differences in synaptic density, mutant dendritic morphologies were not significantly different from wild-type, suggesting that the effect is independent of overall arbour morphology (**Figure S2A-D**). Taken, together these findings imply the need for precise spatial positioning and synaptic transmission during development to establish appropriate synaptic density [53, 57].

### 2.3 Wiring optimisation and synaptic density

A key question is why neurons exhibit a specific synaptic density of 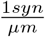. Geometric constraints between neurite cable and volume point to an answer. The scaling relationship between the length of the dendritic cable (*L*) and the volume occupied (*V*) in a circuit is expressed by the scaling law 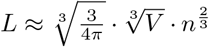, where (*n*) represents the number of synapses attached to the dendrite, assuming dendrites minimise cable length and conduction delays [19, 20] (**Figure 1A**). We compared the predicted dendritic length from this scaling law with the observed values across species, and found that the two align closely to the unity line (**Figure 3A, B**;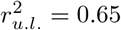), indicating that the scaling law provides sensible predictions across neuronal types and sizes (see **Table 3** for which datasets were used; **Figure S3** for species specific trends; and Methods 3.5 for volume calculations). Residuals were negatively correlated with observed dendritic length (*r* = –0.39, *p* < 0.001), indicating that the model tended to slightly overestimate small dendrites and underestimate large ones. Overall, these results suggest that dendritic arbour scaling is broadly consistent with wiring optimisation theory.

**Table 3.**
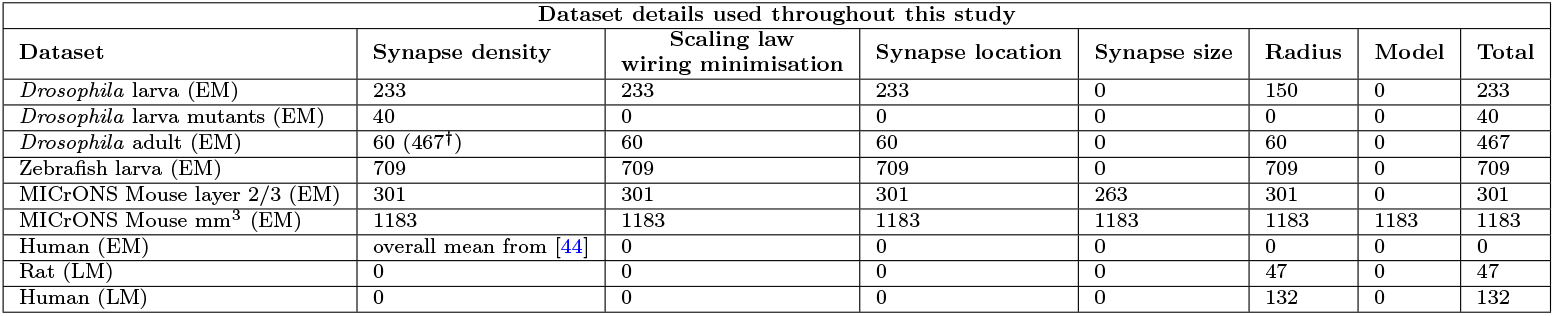
Quantitative summary of number of neurons per dataset used in this study. This table provides a detailed overview of the datasets used in the study, specifying the number of neurons used to compute synapse density, scaling law based on wiring minimisation, synapse location, synapse size, radius, model simulations, and the total count of neurons per dataset. † Indicates the total number of cells used to calculate the postsynaptic reconstruction correlation with synaptic density (**Figure S1 D**).

**Fig. 3.**
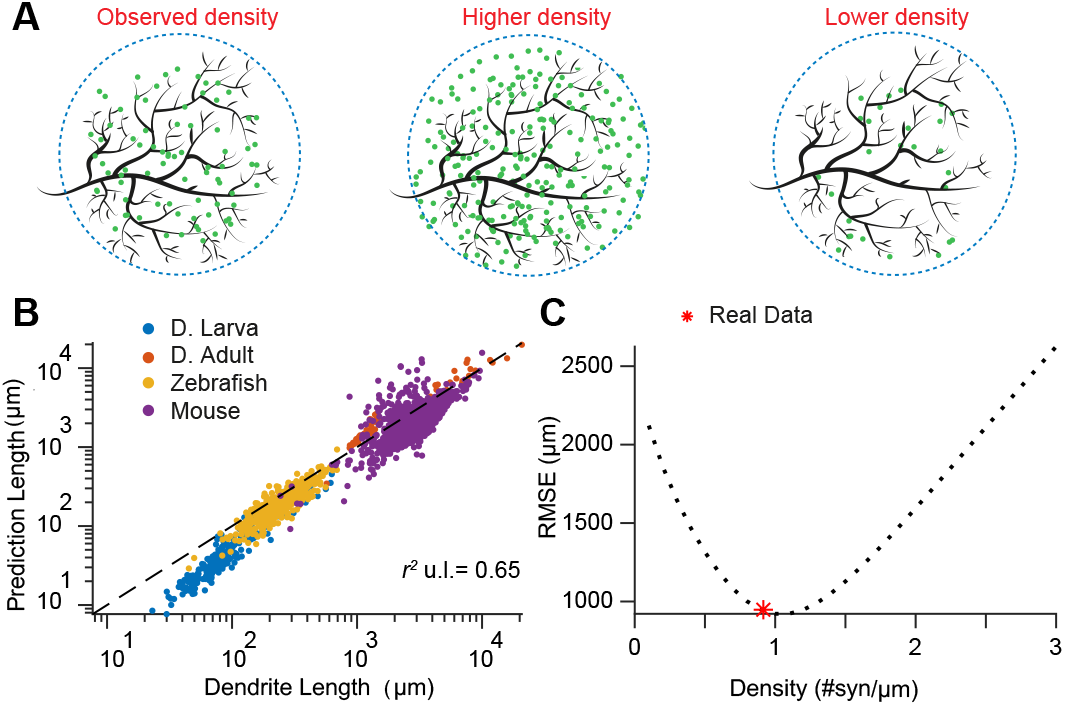
Dendritic structure scaling relationships and synaptic density across species. **A**, Schematic illustrating the impact on synaptic density by varying the number of synapses (green) while keeping dendritic length (black) and volume (blue dashed ellipse) constant. **B**, Shown is the relation between total dendrite length of real dendrites (*L*_*Real*_) and the predicted length (*L*_*Predicted*_) computed using the scaling law for all morphologies of all datasets besides human data (*N* = 2486; log-transformed data; see Table 1, 2). Predicted dendritic length from scaling equation closely matches observed length across all species, with data falling along the unity line (dashed line;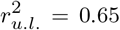; *slope* = 1; fit using linear regression.). Each datapoint is one neuron, with the colour corresponding to species, similar to **Figure 1B. C**, Sensitivity analysis of synaptic density and wiring optimisation. The root mean square error (RMSE) between *L*_Predicted_ and *L*_Real_ is plotted as a function of synaptic density. This analysis systematically varies the synaptic density across all morphologies to evaluate wiring prediction error. This analysis reveals an inflection point around 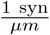. The red star denotes the empirically observed mean synaptic density across datasets.

The scaling law serves as a robust framework for testing how synaptic density relate to wiring costs. We used the scaling relationship to challenge the hypothesis of an invariant synaptic density of 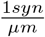 by systematically varying this parameter and calculating the root-mean-square error (RMSE) between the model’s predictions and the observed dendritic lengths (**Figure 3A, C**). The analysis revealed a clear U-shaped curve, with a minimum error at a density of approximately 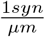, closely matching the average density observed experimentally (**Figure 3C**; see Methods 3.5). At this optimal density, the RMSE was approximately 900 − 1000*μm*. This corresponds to a fractional RMSE of about 40 − 50% to the mean dendritic length across species. Taken together, the close alignment between our sensitivity analysis and measured data, coupled with the sharp increase in error outside this optimal range (up to *RSME* ≈ 2500*μm*, a fractional change of approximately *RSME* ≈ 130 − 140%) supports the hypothesis that the observed 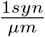 density is shaped by pressure to maintain efficient wiring [58]. However, the remaining error also indicates that additional biological constraints beyond wiring optimisation contribute to dendritic morphology.

### 2.4 Synaptic density correlates weakly with synaptic volume constraints

Next, we sought to understand why synaptic density fluctuates around the mean value of 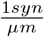. Under the dendritic constancy framework [29], somatic depolarization scales proportionally with synaptic density (*V*_syn_ ∝ *ρ*), implying that the ∼12-fold range observed in our dataset 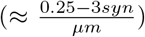 could, in theory, produce a corresponding ∼12-fold difference in voltage response under otherwise constant conditions. Such deviations are predicted to disrupt functional homeostasis, pointing at the importance of identifying their underlying sources.

Insights from cable optimisation theory suggest that reducing conduction delays requires synaptic packing (the volume occupied by synaptic structures) to preserve the optimal 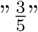 ratio of axon-dendrite volume to total brain tissue [18]. Since we showed that dendritic scaling is consistent with pressure to maintain efficient wiring, we reasoned that optimal packing could regulate the “volume budget” that synapses can occupy in a circuit. However, packing a large number of small synapses, versus a smaller number of large synapses, impacts dendritic synaptic density in opposite ways (**Figure 4A**). Could differences in synaptic size, then, explain the variability in synaptic density around 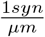?

**Fig. 4.**
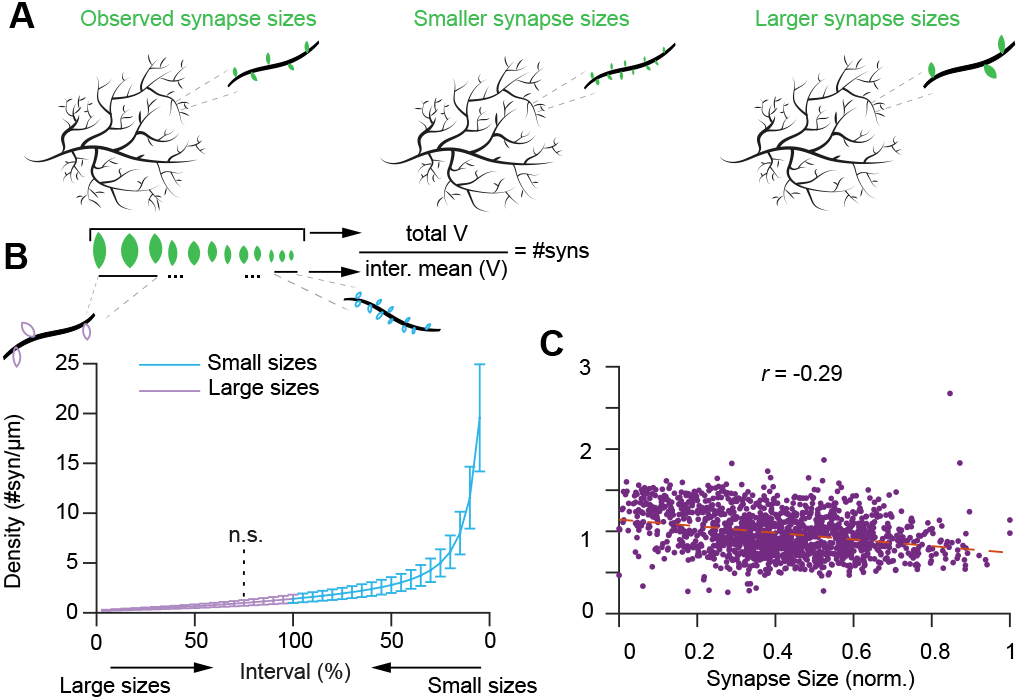
Impact of synaptic size packing on dendritic synaptic density in mouse cells. **A**, The schematic illustrates how different synapse packing arrangements affect synaptic density, without changing the total synaptic volume (shown in green). This effect is observed as synapses are dispersed along a dendritic arbour (black) in the form of numerous smaller synapses or a reduced number of larger synapses. **B**, Schematic (top) and plot (bottom) illustrating how synapse size distribution affects estimated dendritic synaptic density. In the schematic, synapses are sorted by size (green shapes) and grouped into cumulative intervals (lower black bars), starting either from the smallest or largest synapses. For each interval, the total synaptic volume (total V) is converted into a number of synapses using the mean size within that interval (inter. mean (V)). For example, in the top right (blue), total volume is redistributed using smaller synapses, resulting in a higher estimated synapse count and density. In the plot below, the x-axis represents cumulative intervals as percentages, computed in 5% steps starting either from the largest 5% of synapses or the smallest 5%, continuing until all observed synapses are included (100%). The y-axis represents the overall mean synaptic density corresponding to these intervals. Different colours indicate distinct starting points for interval calculation. Error bars indicate standard deviation. **C**, Correlation plot of mean dendritic synapse size (normalised data; *N* = 1446 from [46, 47]; see Methods 3.5) against dendrite synaptic density 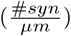 in mouse cells (correlation coefficient of *r* = −0.29, *p <* 0.001). Data points represent individual cells, colours as before. The data are fit with a line of slope −0.4 ± 0.04 (orange dashed line) to assess the relationship between synapse size and synaptic density. Statistical significance was assessed using permutation test (10000 permutations), with Bonferroni correction applied across all interval comparisons to control for multiple testing; “n.s.” denotes non significant. Orange dashed line was fit using linear regression.

To test whether synapse size contributes to variability in synaptic density, we leveraged mouse EM data [46, 47], the only dataset in which synaptic size was measured across most reconstructed morphologies (*>* 94% morphologies; **Table 3**). For each neuron, we first computed the total synaptic volume. We then estimated how many synapses would be needed to fill this same total volume if the synapses were uniformly small or large. To do this, we divided the total volume by the mean synapse size computed within cumulative intervals of the neuron’s synaptic size distribution, starting either from the small or the large synapses. This yielded an estimated synapse count and corresponding synaptic density for each interval (**Figure 4B**; see 3.5).The predicted densities spanned a much broader range than those observed in the real data, with synthetic values ranging from 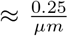 to 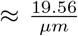, while the observed range was narrower (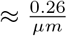to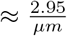). Permutation tests across intervals (Bonferroni-corrected threshold *α* = 0.0013) confirmed that all predicted densities differed significantly from the observed distribution, except for a single interval (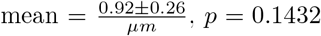; **Figure 4B**).

To better understand the relationship between synapse size and synaptic density, we calculated the correlation between the mean synapse size and the synaptic density for each neuron in the dataset. The results showed a low negative correlation (**Figure 4C**; *r* = −0.29, *p <* 0.001). Taken together, these results indicate that while synapse size and packing constraints can influence density, they alone do not fully explain its variability across neurons.

### 2.5 Synaptic density fluctuations correlate with dendrite thickness to provide voltage response stability

After ruling out packing constraints as the main driver for synaptic density fluctuations, we considered further factors that could explain the variability of synaptic density around the observed mean of 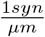. Although noisy developmental processes [53], measurement errors [59], and synaptic plasticity [60, 61] may explain some variance, cell-intrinsic factors are known to strongly impact neuron excitability. These factors include anatomical differences in dendritic radius and variations in the composition of ion channels [62–69]. Interestingly, recent studies have shown that species with larger neurons not only exhibit thicker dendrites but also possess higher membrane ionic conductances, leading to a conserved conductance per unit of dendrite volume that helps maintain stable neuron excitability across scales [26]. Drawing from these findings, we hypothesised that dendritic radius in our dataset may co-vary with synaptic density, with this variability acting as a compensatory mechanism to offset excitability changes arising from structural differences. This relationship could maintain neuronal activity homeostasis and account for the observed variability in synaptic density.

To test this, we leveraged mouse data, for which radii information has been reconstructed across all morphologies, and quantified the correlation between synaptic density and dendrite radius. We found a substantial positive correlation between these features (*r* = 0.67, *p <* 0.001; **Figure 5A**), suggesting that variations in dendritic radius help explain the observed variability of synaptic density around the mean. This correlation could also account for previous reports of slightly higher values of synaptic density in mouse data than 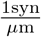 [44, 70].

**Fig. 5.**
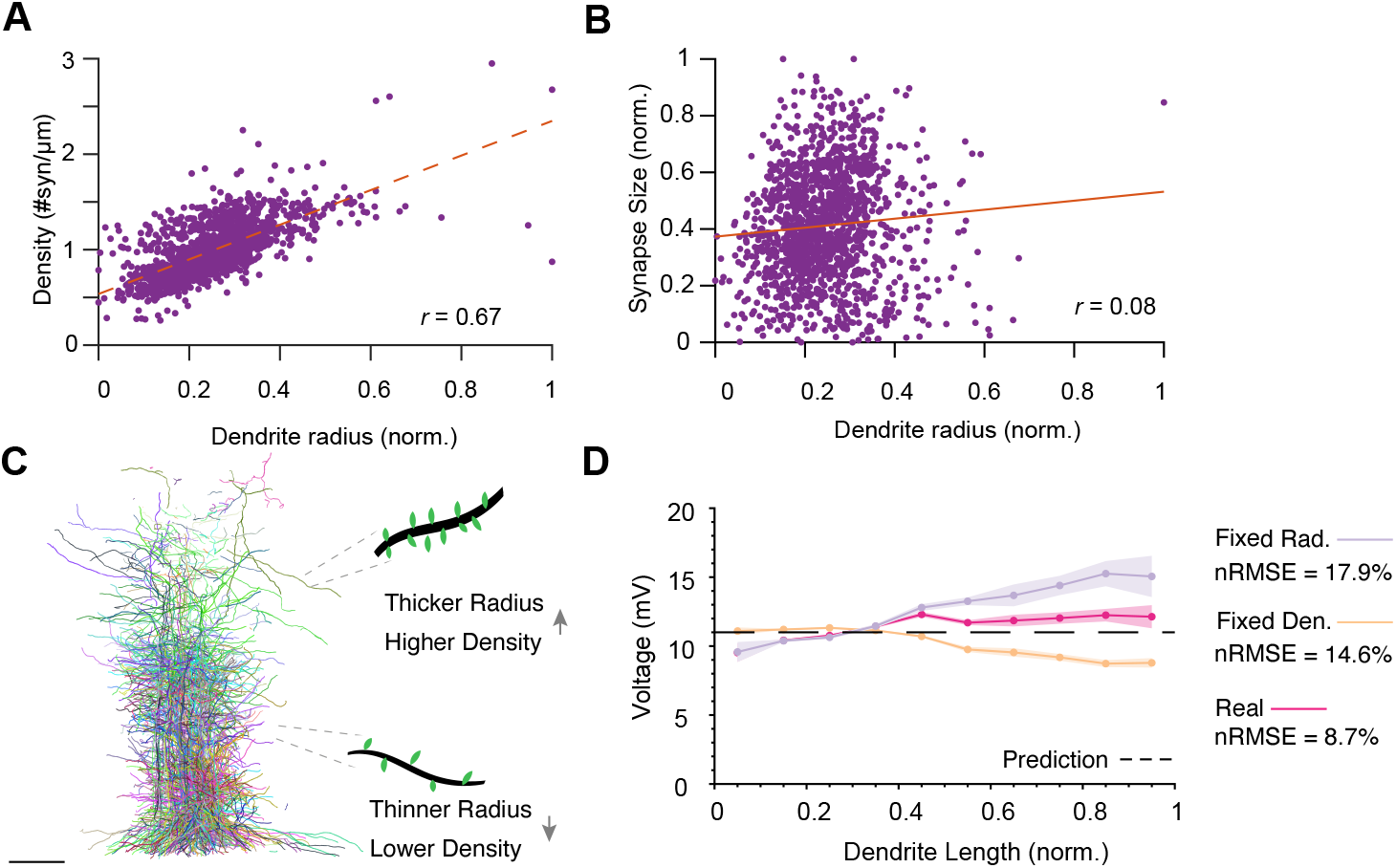
Radius-synaptic density correlation stabilizes voltage response in steady-state biophysical models. **A**, Correlation plot of normalised dendrite radius (*N* = 1484; see Methods 3.5) against synaptic density 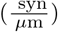 in mouse cells. The correlation coefficient is *r* = 0.67 (*p <* 0.001). Data points represent individual cells, and colours match prior panels. The data are fit with a line of slope 1.81 ± 0.05 (orange dashed line), to assess the relationship between dendritic radius and synaptic density. **B**, Correlation plot of normalised dendrite radius (*N* = 1446; see Methods 3.5) against normalised synaptic size in mouse cells. The correlation coefficient is *r* = 0.08 (*p <* 0.01). Data points represent individual cells, and colours match prior panels. The data are fit with a line of slope 0.16 ± 0.05 (orange line), to assess the relationship between dendritic radius and synaptic size. **C**, Shown are a random subset of neuronal morphologies used to constrain the biophysical models. Right, schematic of the proposed compensatory mechanism maintaining neuronal excitability in mouse data. Top: a thicker dendrite adjusts synaptic density to offset higher input conductance, due to the increased membrane surface area. Bottom: the opposite for a thinner dendrite. Scale bar 100*μm* **D**, Average voltage responses of modelled morphologies (*N* = 1186) plotted against normalised length under three conditions: mangenta, real diameters and synaptic densities; light purple, real synaptic densities but average fixed diameters across all morphologies; yellow, real diameters but average fixed synaptic densities. Corresponding nRMSE values are provided for each condition. Shaded areas represent standard error of the mean (SEM). Black line show predictions from the dendritic constancy equation. Target voltage was set to *V*_*syn*_ = 11*mV*, to match spike threshold of mouse pyramidal neurons following [29]. Correlation coefficients (*r*) are calculated using Pearson correlation; *p* values are obtained from permutation tests (10000 permutations), and linear regression lines are fit where applicable.

Recent studies show that the area of postsynaptic densities (PSDs) serves as a proxy for synaptic strength, with input synaptic current increasing linearly with PSD area [71]. To test whether synapse size could confound the hypothesised compensatory relationship between synaptic density and dendrite radius, we examined the correlation between synaptic size (which is highly proportional to PSD area in mouse [47]) and local dendritic radius across all mouse morphologies for which both features were measured. We found a weak relationship (*r* = 0.08; *p <* 0.001; **Figure 5B**), indicating that synapse size does not systematically vary with dendritic diameter. However, since synaptic size and density are modestly negatively correlated (**Figure 4C**; *r* = − 0.29), we cannot exclude a small compensatory role for synaptic size. Nonetheless, these findings support our conclusion that it is not the primary driver. We therefore focus on dendritic radius as the primary structural variable influencing synaptic input integration.

To test whether the observed correlation between dendritic radius and synaptic density can stabilise voltage responses, we developed passive single-cell models based on mouse MICrONS mm^3^ reconstructions, with distributed synaptic inputs placed along their dendritic trees (**Figure 5C**; **Table 1, 3**; see Methods 3.5). Biophysical parameters were constrained using published experimental measurements from cortical pyramidal neurons [29]. We compared the simulated somatic voltage responses of these models to analytical predictions from the dendritic constancy framework, which posits that steady-state voltage responses remain invariant to neuron size when synaptic density is fixed. Specifically, the expected voltage at the soma is given by (*V*_syn_), which can be approximated as 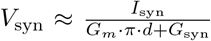, where *I*_syn_ is the total synaptic input current representing the sum of currents from distributed synaptic inputs, *G*_*m*_ is the membrane conductance reflecting the electrical properties of the dendritic membrane, *d* is the average dendritic tree diameter, and *G*_syn_ is the total synaptic conductance. All terms in the equation were constrained using the same values used in the biophysical models to ensure consistent comparison under physiologically realistic conditions.

To test the effect of realistic dendritic radius and synaptic density fluctuations on voltage responses, we simulated three conditions: (1) using cell-specific dendritic diameters with fixed average synaptic density across the dataset, (2) using cell-specific synaptic densities with fixed average dendritic diameters, and (3) using the measured values of both radius and density for each neuron (**Figure 5D**; see Methods 3.5). When synaptic density was fixed, cells with larger diameters showed attenuated responses due to increased membrane conductance (*nRMSE* = 14.6%). When diameter was fixed and synaptic density varied, longer cells produced larger voltage responses (*nRMSE* = 17.9%). However, when real radius and density values were paired, deviations were minimized (*nRMSE* = 8.7%). In all cases, nRMSE was computed relative to the theoretical voltage predicted by the dendritic constancy equation. These findings support the hypothesis that co-variation between dendritic radius and synaptic density serves to maintain stable neuronal excitability.

### 2.6 Simulated co-scaling of synaptic size and dendritic radius supports excitability stability

Although our ultrastructural datasets lack sufficient dendritic radius information to test the above hypothesis comprehensively across species, we acknowledge that this strategy can become self-defeating across scales. Given that synaptic density converges to approximately 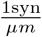 across species, and that dendritic radius increases with brain size to reduce conduction delays [72], it follows that synaptic density cannot scale up indefinitely as brains enlarge. This underscores the need for additional compensatory mechanisms to maintain stable voltage responses. What alternative factors could compensate for higher input conductance due to thicker cable calibre?

According to the dendritic constancy equation, for a given synaptic density and assuming membrane conductance and input current remain constant, an increase in dendritic radius reduces voltage responses. This effect could, in principle, be offset by a proportional increase in synaptic input. However, since synaptic density remains largely invariant across species, this compensation cannot be achieved by adding more synapses. Instead, synaptic strength at the level of individual synapses would need to increase. Given that synaptic size correlates with synaptic strength [71, 73], we reasoned that PSD area might increase with dendritic radius across species to offset the higher input conductance introduced by thicker dendrites (**Figure 6A**). To explore this, we combined PSD and radii data from our own measurements with data from the literature to complement missing species-specific datasets (**Table 3, 4**; see Methods 3.5). Interestingly, we found that the mean size of the synapse, measured by PSD area, shows a significant strong correlation with the average dendritic radius between species (*r* = 0.88, *p <* 0.018; **Figure 6B**). This correlation supports the hypothesis that interspecies differences in synaptic strength could partially compensate for changes in excitability resulting from radii scaling.

**Table 4.** Details of species, brain regions and quantification of postsynaptic density area. The table summarizes the postsynaptic density (PSD) characteristics for various species, including *Drosophila* (larva and adult), rat, mouse, and human. For each species, the specific brain region studied, sample size, and mean area of the PSD in square nanometres (*nm*^2^) are provided. For human data, “AS” denotes asymmetrical synapse, and “SS” symmetrical synapse. References for the data sources are included.

| Postsynaptic density details derived from electronic microscopy |  |  |  |  |
| --- | --- | --- | --- | --- |
| Species | Brain region | Sample size | Mean area ($nm^2$) | Neurotransmitter type |
| *Drosophila* larva [121] | Antenal lobe and ventral nerve chord | 540 | 14527 | Excitatory |
| *Drosophila* adult [122] | Mushroom Body | 100 | 26001 | Excitatory |
| Rat [97] | Hippocampus | 287 | 50800 | Excitatory |
| Mouse [71] | Neocortex layer 2/3 | 15 | 74667 | Excitatory |
| Human [123] | Entorhinal cortex (AS) | 8794 | 118492 | Excitatory |
| Human [123] | Entorhinal cortex (SS) | 550 | 70501 | Inhibitory |

**Fig. 6.**
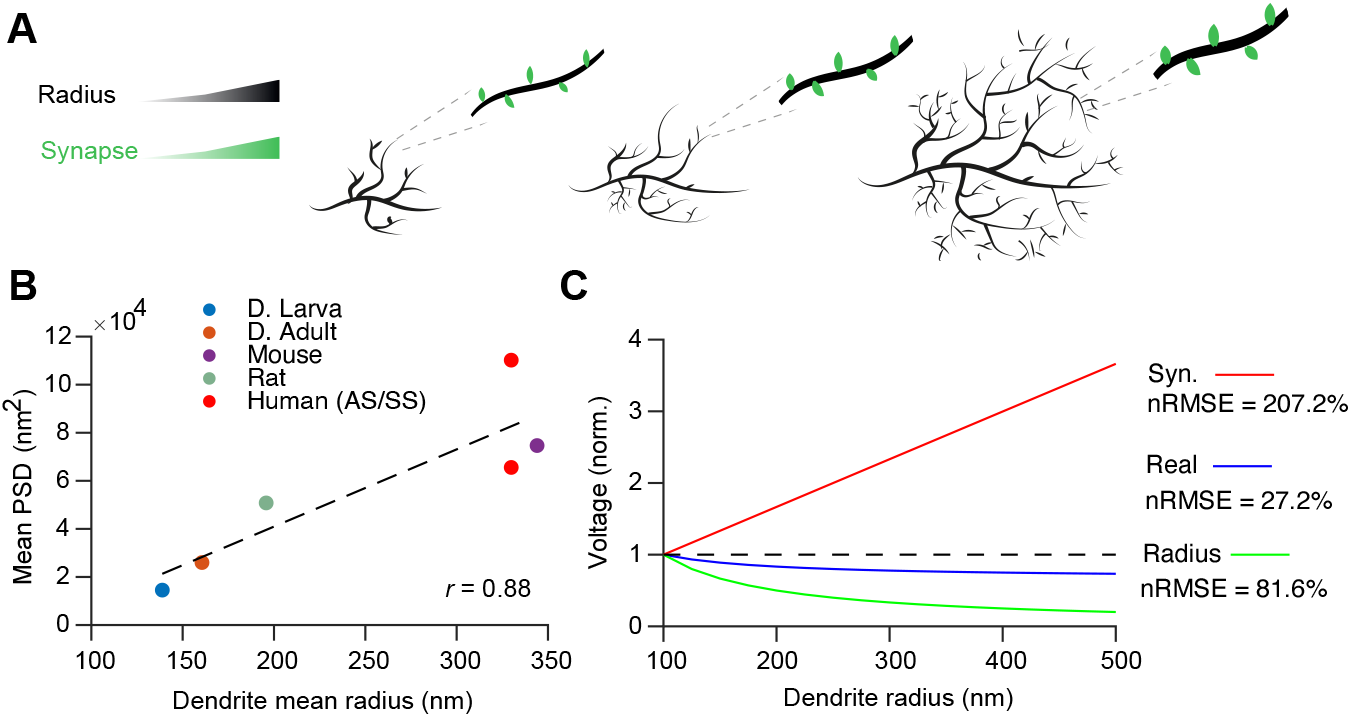
Relationship between dendritic radius and synaptic size across species. **A**, Schematic of the hypothesised compensatory relationship for maintaining neuronal excitability across species. As dendrites (black) increase in size, synapses (green) also increase in size to offset higher input conductance and stabilise the voltage response. **B**, Correlation plot between the mean radius of dendrites (*N* = 3309; **Table 3**; Methods 3.5) and the size of synapses (*N* = 10286; **Table 4**; Methods 3.5), measured by the area of postsynaptic densities (PSDs), across different species (correlation coefficient of *r* = 0.88, *p* = 0.018). Each color represents a different species. For human data, “AS” is asymmetric synapses (larger PSD value) and “SS” is symmetric synapse (smaller PSD value). The data are fit with a line of slope 321.25 ± 87.82 (black dashed line) to assess the relationship between dendritic radius and synaptic size (PSD). **C**, Normalised voltage responses plotted against dendritic radius (range = 100–500*nm*) under three conditions. The dashed line represents the initial voltage response at the beginning of the range, which was used to normalise all other responses. We quantified deviations from the initial response using nRMSE. Blue: dendritic radius, input current, and synaptic conductance scale linearly. Red: input current and synaptic conductance scale, but radius remains fixed at 100 nm. Green: radius scales while input current and synaptic conductance remain constant. Correlation coefficients (*r*) were obtained using Pearson correlation, and *p* values obtained from a permutation test (10000 permutations); lines were fit using linear regression.

To evaluate whether synaptic strength could compensate for changes in dendritic radius, we used the dendritic constancy equation to predict neuronal voltage responses across three parameterisations of dendritic radii 100–500nm, covering the full range of values in our dataset (**Figure 6B**; see Methods 3.5). First, we scaled both synaptic input current and synaptic conductances linearly with dendritic radius, based on the correlation between radius and synaptic size in **Figure 6B**. Second, we scaled only the dendritic radius while fixing input current and synaptic conductance at the lowest predicted value in the range. Third, we scaled synaptic parameters while keeping the radius at the lowest value. We found that when input current and synaptic conductance were kept constant, increased dendritic radius led to decreased voltage responses and deviations from initial values (**Figure 6C**; nRMSE = 81.6%; nRMSE calculated against the initial voltage value). Conversely, scaling input current and synaptic conductance with a constant radius resulted in significant increases in voltage responses, with higher deviations (**Figure 6C**; nRMSE = 207.2%). In contrast, co-scaling dendritic radius and synaptic parameters yielded responses closely matching initial values, with minimal deviation (**Figure 5C**; nRMSE = 27.2%). These findings suggest that joint scaling of dendritic radius and synaptic strength may counterbalance their opposing effects, maintaining stable voltage responses across the observed range of dendritic radii.

## 3 Discussion

The mechanisms and constraints that dictate the scaling of nervous systems are still poorly understood. In this study, we showed that synaptic density is remarkably constant around 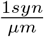 across different neuronal types and species. We found that synaptic density is shaped by spatial positioning and synaptic transmission during early development. Furthermore, we provided experimental evidence that validates a longstanding morphological scaling law derived from optimal wiring principles [19]. Using these quantitative relationships, we predict that synaptic density specifically converges to 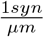 due to pressure to optimise cable resources. Finally, our simulations make predictions that synaptic density of 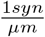 synergises with other morphological features to maintain voltage responses stable after synaptic input integration. Our results suggest that is an optimal set point for brain architecture and computational reliability of neurons, while minimising resources as the nervous system scales across evolution [10].

### 3.1 Limitations of EM data in this study

A central assumption of this study is that dendritic synaptic density remains stable after early development and is conserved across species. One limitation of EM-based connectomics is that it captures only static snapshots from a small number of specimens, raising questions about generalizability [74]. Sample sizes are often small, and reconstruction quality can vary across datasets [75]. To mitigate these issues, we restricted our analysis to neurons with near-complete dendritic reconstructions and applied only minimal corrections, grounded in known technical artifacts such as segmentation errors and incomplete proofreading (Methods 3.5). This conservative approach meant excluding several available datasets and species, reflecting a trade-off where completeness of analysis was prioritized over increasing the number of datasets [34, 37, 76]. Thus, anatomical sampling is uneven across species, with the number of adult *Drosophila* reconstructions being on the low end, and zebrafish data limited to the olfactory bulb [41, 77]. These factors could bias density estimates, distribution spread, or obscure subtype-specific variability. Nonetheless, the convergence toward approximately 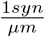 was consistently observed, and our two most densely reconstructed datasets, *Drosophila* larva [33] and mouse [46, 47], show no significant difference despite their vast phylogenetic and anatomical differences.

Can we trust that the variance around the mean synaptic density reflects biological variability rather than technical noise [78]? Within our curated dataset, we believe so. The variability is not random but appears to be systematically compensated by co-variation with dendritic radius, supporting the stability of voltage responses. One limitation, however, stems from the lack of standardization in EM data mapping [75]. While some datasets provide detailed metrics—such as synapse size, dendritic radius, and full morphologies—others include only a subset of these features. For example, detailed measurements of synaptic size and dendritic radius, which we found to correlate with synaptic density in mouse, were not available in other species. As a result, we cannot determine whether this compensatory mechanism generalizes broadly. Future cross-species datasets with consistent morphometric measurements are necessary to uncover additional compensatory strategies.

Beyond anatomical factors, species- and cell-type-specific variations in ion channel composition significantly impact neuronal excitability [66, 67, 79, 80]. However, electrophysiological features aren’t captured well by anatomical data alone. Thus, in our EM-constrained modelling framework, we assumed homogeneous passive electrical properties across all cells, and only considered excitatory synaptic inputs. This simplification allowed us to isolate the impact of morphological features, namely, dendritic radius and synaptic density, on voltage response. While this approach is appropriate for hypothesis generation, it necessarily limits the physiological realism of the model. Although electrophysiological variables are outside the scope of the anatomical datasets analysed in this study, they could contribute to computational specificity and influence the degree of voltage response homeostasis achieved by synaptic density invariance [29, 81, 82].

Recent work, however, suggests an exciting alternative view, suggesting that ion channel diversity instead provides neurons with flexibility and robustness to maintain target excitability [65, 68]. Theoretical models support this notion, demonstrating that neurons in diverse species can maintain stable physiological properties despite substantial size differences by regulating multiple conductances [69]. Additionally, evolutionary evidence highlights that different species adapt ion channel conductances in conserved cell-types, effectively compensating for anatomical differences to maintain their functional output. [26]. This suggests a biophysical mechanism that regulates intrinsic conductivity, contributing to the stability of neuronal activity despite variations in scale and connectivity across cell-types and species.

In conclusion, these limitations reflect the current state of available EM datasets. Nonetheless, the convergence observed across our data and modelling provides strong evidence that synaptic density is a conserved feature likely to support the stability of neuronal excitability.

### 3.2 Why a synaptic density of 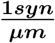?

To gain insight into why synaptic density converges specifically to 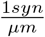 across species, we adopted a normative perspective, asking whether this value represents an evolutionary optimum for wiring efficiency. Dendritic constancy does not prescribe an explicit density value—any fixed density would, in principle, suffice for maintaining voltage response stability. Thus, we hypothesized that synaptic density of 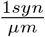 may instead reflect pressures from wiring optimization [18, 19].

To test our hypothesis, we applied a scaling law that predicts a 2/3 power law relation between dendritic arbour structure under assumptions of cable minimization [19]. Residuals from the scaling model showed a modest negative correlation with dendritic length, indicating a tendency to overestimate small arbours and underestimate large ones. However, these deviations align with known species-specific strategies. For example, small quasi-planar arbours in *Drosophila* larvae may fill space more efficiently than predicted by the 3D 2/3 power law, instead following a shallower 1/2 exponent appropriate for planar geometries [19–21]. Conversely, large neurons often increase dendritic length to create more direct paths from synapses to the soma in order to reduce conduction delays [83]. This strategy is known to introduce a vertical offset in the scaling relationship [19]. Despite these boundary-specific deviations, the underlying scaling trend was maintained, suggesting that the convergence of synaptic density to 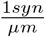 is not arbitrary, but reflects a balance between morphological and functional constraints. Nonetheless, the remaining error in our sensitivity analysis also suggests that additional constraints, such as metabolic efficiency [84], protein transport [85], and synaptic plasticity [60, 61], may further shape synapse allocation in a cell-type- or context-specific manner.

### 3.3 Synaptic density invariance in scaling nervous systems function

What are the functional implications of synaptic density invariance across neuronal types and species in scaling nervous systems [82, 86]? The dendritic constancy principle proposes that voltage response stability depends primarily on synaptic density regardless of cell-type morphology and connectivity [29].

In particular, for a given synaptic density, if the average dendritic radius and membrane conductivity are consistent across neurons, the effect becomes more significant. Neurons then functionally approximate point-neurons (excluding axo-axonic synapses), with dendritic morphology-related effects on synaptic integration substantially reduced [29]. Although dendritic radii and membrane conductivity varies across species and cell-types [72], our analysis suggests that dendritic radius increases with synaptic density across mouse cell-types and with synaptic size across species (**Figure 5, 6**). These co-variations, combined with compensatory mechanisms such as ion channel conductance adjustments, can mitigate the effects of increasing cell size to preserve neuronal excitability [26].

This “tug-of-war” dynamic between structural and biophysical factors may not strictly adhere to the “dendritic radius and membrane conductivity constancy” constraints, but could still produce equivalent functional outcomes [87]. We speculate that the interplay between synaptic density invariance and compensatory mechanisms enables neurons from the same cell-type to preserve their excitability [29]. Such morphology-independent neurons could endow networks with the capacity to maintain functional robustness regardless of brain size. Further work is needed to determine the extent to which these strategies apply universally across species, circuit architectures, and evolutionary contexts.

To clarify, we are not proposing a normative framework in which neurons operate strictly as point-like integrators. A corollary of the dendritic constancy principle suggests that synaptic density invariance creates a passive backbone for the conservation of excitability, that preserves the ability to modulate spike timing through various factors [29]. These factors include ion channel composition, intrinsic cellular properties, and morphology, which can generate cell-type specific spike-time-dependent computations [23, 29]. Stable excitability is particularly important for reliable computations that depend on spike timing, as they provide a consistent background against which temporal variations can be effectively encoded [23, 88, 89]. Thus, synaptic density invariance may play a critical role in supporting cell-type-specific modulation of spike timing [23, 29, 60, 81, 90, 91].

### 3.4 Implications for interpreting and modelling connectomes

Let us consider how synaptic density invariance can contribute to the practical study of connectomes. For example, reducing cell-type specific morphologies can significantly lower the degrees of freedom in brain-scale modelling [92, 93]. Building on this insight, synaptic density invariance and its implications offer a biologically grounded framework to reduce dendrites into point representations; a simplification that retrospectively supports published modelling studies of *Drosophila* and mouse connectomes, where modellers use deep neural networks or leaky integrate-and-fire models to simulate and predict spiking behaviour within connectome-constrained networks. [92–96].

Relatedly, dendritic constancy may also help in interpreting synaptic connectivity, that is the “edges”, in connectomes. A naive approach assumes that the more synapses that connect two neurons, the stronger that connection must be. However, the absolute numbers of synapses vary between individuals, brain regions, and even within circuits [33, 34, 36, 37, 47, 97], and this variability complicates interpreting connectivity patterns in connectomics datasets [98–101]. Dendritic constancy predicts that, for a given density, postsynaptic voltage responses scale with the proportion of active synapses relative to the total synaptic count on the postsynaptic neuron [29, 102]. This voltage response to relative connectivity mapping is independent of neuron size and overall connectivity, effectively normalising synaptic weights [102–104].

Interestingly, recent EM circuit mappings in *Drosophila* reveal that the relative axo-dendritic synaptic connectivity between neuron partners, in contrast to the absolute number, are conserved between individuals, developmental stages, hemispheres and sexually dimorphic circuits [48, 49, 54, 100, 101, 105–107]. In agreement with this idea, changes in relative connectivity, rather than absolute synapse numbers, caused by experimentally induced developmental perturbations lead to functional and behavioural phenotypes [56], as can relative connectivity changes observed across evolution [108, 109]. In our accompanying paper, we show that maintaining synaptic density and relative connectivity across development, can preserve consistent postsynaptic functional responses even as neurons grow fivefold [110]. Taken together, these findings support interpreting connectivity through the lens of input fractions, suggesting that future comparative studies should focus on relative, percentual connectivity rather than absolute synaptic counts.

### 3.5 Conclusion

Our findings inform an ongoing debate in neuroscience. Dendritic complexity is crucial for understanding brain function [82, 86, 111, 112], however, both simplified models of neural circuits and machine learning research have achieved remarkable success with networks of point neurons that abstract away cellular detail [92, 113, 114]. Our data and analysis support the use of point neuron representations for dendrites as useful abstractions of what nature – across species and scales – converged on to maintain stable neuronal responses; axons, though, with their abundant axo-axonic synapses [33, 47], lay beyond the scope of our analysis. As connectomics advances, and the number of structural maps outpace our functional understanding of a given nervous system, principles like synaptic density invariance will be useful to simplify and interpret connectomes.

## Acknowledgments

We thank Adrian Wanner and Rainer W Friedrich, Casey Schneider-Mizzel, and Sahil Loomba for sharing the zebrafish, mouse and human datasets, respectively. We thank Tom Bartol for sharing the rat postsynaptic density areas dataset; and Sergio Plaza Alonso, Lidia Alonso-Naclares and Javier Defelipe for sharing the human postsynaptic density areas dataset. We also thank Daniel Han, Philipp Schelegel, Dániel Barabási and Benjanmin Piazza for support in handling various connectomics datasets. The authors thank Marc Corrales and Simon Laughlin for helpful comments in the manuscript. Finally, we thank Ivana Henry for helping design some sketches. We also acknowledge Ingo Fritz, Ricardo Chirif Molina and Feiyu Wang for help collecting postsynaptic density data.

Supported by Wellcome Trust Investigator Award 205038/Z/16/Z to A.C., and core funding from the MRC LMB. The project was also supported by a Daimler-Benz Grant (A.F.C.).

## Author contributions

André Ferreira Castro, Conceptualisation, Resources, Data curation, Investigation, Formal analysis, Methodology, Writing; Albert Cardona, Conceptualization, Resources, Data curation, Methodology, Writing.

## Data and code declaration

Availability of data and materials: all data and code are available at https://drive.google.com/drive/folders/11FUuRuDHUt5q_IM9wfKY-Y5PxDcH0nNW?usp=sharing.

## Methods

### Electron microscopy data and morphologies

Morphologies were obtained from five electron microscopy volumes encompassing the central or peripheral nervous systems of four distinct species, including one first instar *Drosophila melanogaster* larva [33, 106, 115], one adult *Drosophila melanogaster* [41], one larval Zebrafish (*Danio rerio*; [45]), and finally two adult mice (*Mus musculus*; [46, 47]). These datasets were selected because they were densely reconstructed, annotated and proofread by experts.

From the larval insect datasets, neurons were selected based on the reconstruction level of the corresponding cell types (**Table 1**). Although the larval complete connectome has been released, only a few brain regions have been fully reconstructed and have all anatomical synaptic sites annotated [33]. To maintain consistency across the brains of larval and adult flies, we pooled the same cell types from the Hemibrain dataset. However, the postsynaptic reconstruction level for some of these cell types were low, which were highly correlated with synaptic density (*r* = 0.61, *p <* 0.001, **Figure S1D**). To circumvent this issue and avoid bias in our analysis, we discarded cell types with postsynaptic reconstruction levels below 80%. To increase the sample size, we searched for other regions of the brain with adequate levels of reconstruction.

This search yielded another brain region that satisfied this constraint, the ellipsoid body [77]. For the case of the ellipsoid neurons, we included all neurons marked as fully traced.

Additional data was analysed spanning 4 volumes of electron microscopy (**Table 1**), including two *Drosophila melanogaster* larva mutants (*FraRobo* and tetanus toxin light chain *TNT* ; [56]), and data from two cerebral cortices of humans [44]. From [44], human neuron morphologies were not used directly because the data was not released; instead, only estimates of synaptic counts in dendritic trees and the total length of dendritic trees of the available cell types were used to compute the general dendritic synaptic density.

### Measurement of dendritic radius

To measure dendritic radii, we used data from the aforementioned *Drosophila* and mouse EM datasets, and additional light microscopy (LM) datasets pooled from the neuromorpho.org database (see **Table 1**). For the EM datasets, raddi data were readily available from three EM volumes of two distinct species, adult *Drosophila melanogaster* [41], and two adult mouse (*Mus musculus*; [46, 47]). However, the radius data from the adult *Drosophila* EM datasets were not proofread by expert annotators; therefore, they were excluded from finer analyses, such as comparing radii with synaptic density. Instead, they were used in coarser analyses, such as examining interspecies comparisons and PSD size variations across species.

In the case of the *Drosophila* larva EM dataset [33], the radius information was missing. To address this, we used a custom tool in CATMAID [116] to manually trace the radius of 20 full dendrites of Kenyon cells (KC) from the mushroom body (MB). To increase the representation of our dataset, we first computed the mean radius of the nodes for each Strahler order of manually traced KC dendrites [117, 118]. Next, we determined the Strahler number for all other KC dendritic nodes in both MBs (left and right). Using the mean radius value corresponding to each Strahler order, we assigned these values to each node of the non-traced KCs dendrites. We then calculated the mean radius value on the entire KCs dataset (*N* = 150). Thus, ensuring a consistent and representative measurement across the dataset.

From the light microscopy datasets, we included data from human [119] and rat [120] to match the available ultrastructural postsynaptic density (PSD) dataset (see **Table 3, 4**). For the rat dataset, it was possible to match LM morphologies with areas of the brain where PSDs were quantified. However, for human data, it was not possible to match the LM morphologies to the specific brain regions where the PSDs were obtained. Therefore, we sampled all available morphologies from the NeuroMorpho database to avoid sampling biases. Similarly to EM datasets, only data annotated with complete 3D dendritic reconstruction status were used to compute the dendritic radius.

### Measurement of postsynaptic density

Postsynaptic density (PSD) areas were obtained from various EM volumes that encompass the synapses of the central or peripheral nervous systems of four distinct species (Table 4), including *Drosophila* (larva [121] and adult [122]), mouse [71], rat [97] and human [123]. These data sets were selected because they provide ultrastructural resolution data on PSD areas, which is a strong anatomical proxy of synaptic strength [71, 73].

For the adult *Drosophila* dataset, we collected PSD areas following the procedure described by [121] for synapses of interest from the FAFB dataset [122]. Shortly, using a CATMAID’s Python interface, we extracted the locations of synaptic contacts between neurons of interest. With BigCAT (https://github.com/saalfeldlab/bigcat), a BigDataViewer-based volumetric annotation tool, we annotated the postsynaptic membrane across EM slices. The synaptic surface area is represented by all visible membrane sections, measured in *nm*^2^. The total area of a synapse is approximated by summing these areas for a single contact across z sections.

### Neuronal morphologies and skeletonisation

Electronic microscopy neuronal skeletons for adult and larval *Drosophila* were obtained from virtualflybrain.com using custom Python and R scripts. The mouse arbour morphologies were obtained by downloading them from www.MICrONS-explorer.org or requesting them directly from the authors of the original studies [47]. In the case of larval zebrafish, skeletons were derived from the meshes of the corresponding neuron morphologies provided by the authors of the original study [45]. These meshes were imported into the Navis environment (https://github.com/navis-org/navis), where we extracted the spatial coordinates of each skeleton node along with soma identifiers for further analysis. For the case of LM data, neuronal skeletons were obtained from the neuromorpho.org database [124].

Next, all skeletons were imported into the TREES Toolbox (MATLAB; https://www.treestoolbox.org) for quality control, in a two-step process. Initially, we used specific inbuilt functions in TREES Toolbox to verify data integrity, followed by a visual inspection by two expert annotators. In some cells, the ambiguity in the location of the soma within the representations of the skeleton was resolved through consensus of two expert annotators, who used cell type-specific anatomical features and radius information to estimate the correct location of the soma. Neurons with persistent ambiguity after proofreading, mainly due to artefacts, were excluded from the analysis, generating a total number of skeletons *N* = 3112, for all species (**Table 1**).

In the larval zebrafish dataset, we noted that some skeletons presented cycles in their graph representation, a common error when segmentation-based proofreading is used for skeletonisation. This error typically arises due to the addition of false side branches in voluminous neuronal parts such as the soma, or due to image artefacts or masking issues. To address this, we first resampled the trees to a distance of 1*μm* and then used a previously described minimum-spanning tree (MST)-based algorithm to reconnect the nodes to skeletons [83]. This effectively maintained the original tree’s topology and geometry while eliminating cycles from the graph structure.

Finally, all EM-derived data, namely skeletons, synapse, and soma location data, were imported into Navis (Python, https://github.com/navis-org/navis), Natverse (R, [125]), or TREES Toolbox (mathworks.com) for subsequent analysis, and were rescaled to 1×1×1 *μm*. Custom Python, R, and MATLAB scripts were used analyse skeletonstatistics, performed within the aforementioned environments. All analysis scripts and files related to neuronal morphology and synapse locations will be available upon publication.

### Distinguishing axons and dendritic compartments in neuronal trees

To accurately test the predictions from the dendritic synaptic density, synaptic packing, and wiring minimisation scaling law, the arbours of all available neurons were split into axon and dendrite compartments. Then the dendritic sub-arbour was selected for further analysis, discarding the axonic compartment. For the cases of the first instar Drosophila larva [33, 106, 115], and adult mouse datasets [46, 47], existing expert annotations were used to split the neurons.

For ellipsoid body neurons, we computed the segregation index (SI), a measure of neuronal polarisation as established by [59]. An SI of 1 indicates total segregation of inputs and outputs into dendrites and axon, while an SI of 0 indicates a homogeneous distribution. For neurons with a low segregation index (SI *<* 0.05), we considered the entire arbour as a dendritic tree as described in [59]. Therefore, we treated these unsegregated neurons as purely dendritic, only removing the main trunk (linker segment) that stems from the soma of the rest of the analysis.

The remaining neurons from adult *Drosophila*, and larval zebrafish, were divided into axonic and dendritic compartments using a newly developed strategy that combines K-means clustering algorithms to find the nodes of the arbour that belong to the dendritic or axonic compartments, followed by consensus validation by 2 expert annotators (**Figure S4**).

### Axon-dendrite split in the adult *Drosophila*

*Drosophila* neurons exhibit a diverse range of anatomical structures, reflecting their various functions within the insect’s nervous system. However, a typical polarised neuron in *Drosophila* consists of a cell body (soma), from which an extended main branch connects a dendritic tree upstream, and axonal arbours downstream ([33, 41]; **Figure S4A**). Our new algorithmic approach takes advantage of those inherent differences in the structural characteristics of fly axons and dendrites to effectively separate them, applying the K-means clustering method to partition the nodes of neuronal trees into axonal and dendritic compartments (**Figure S4A**, step 1). The K-means clustering algorithm is a well-established technique used for partitioning a dataset into K distinct, non-overlapping subsets (clusters). The algorithm identifies cluster centres (means) that minimise the sum of squares within the cluster (i.e., the Euclidean distance from each data point in the cluster to the respective mean).

Our strategy begins by representing each node in the neuronal tree as a 3D data point. Each dimension corresponds to a distinct coordinate of the node in space. Next, the K-means algorithm is applied to this 3-D dataset, with K set to 2, representing the two primary compartments of the neuron: the axon and the dendritic tree. The algorithm iteratively assigns each node to the cluster (compartment) whose mean is closest, then recomputes the means. This process is repeated until the assignment of nodes to clusters no longer changes. The choice of K as 2 is rooted in the biological structure of the neuron. Although there may be substructures within the axon and dendritic compartments that could be further subdivided, our aim was to initially segregate these two primary compartments (**Figure S4A**, step 2). Following the initial clustering that identifies two distinct compartments using our strategy, we make an anatomically informed assumption to further segregate the dendritic compartment. Given that the dendritic compartment is typically closer to the soma, we consider the cluster containing the soma node as the dendritic compartment in our first-pass analysis. Subsequently, we further parse the dendritic compartment, removing the main branch nodes contained in the initial dendritic compartment. We compute the longest path length from the soma to the end of the arbour to identify the main branch, a criterion commonly used in computational neuroanatomy to identify the main branch of a dendritic tree [20]. This is based on understanding that the main branch of a neuron does not constitute a significant part of the dendritic arbour and is typically longer than other branches (**Figure S4A**, step 3). The dendritic compartment is then redefined as the set of nodes found in the initial dendritic compartment minus the main branch. This method allows us to further refine our anatomical segregation of the neuron, allowing more detailed and granular analyses of the neuronal structure (**Figure S4A**, step 4).

### Axon-dendrite split in larva zebrafish

In the case of zebrafish mitral cells, we slightly adapted our neuronal compartment K-mean clustering strategy to account for the unique morphology of these neurons (**Figure S4B**, step 1). The typical structure of an adult zebrafish mitral cell is characterised by a soma, from where a single primary dendrite terminating in dendritic tufts near the cell body, and a single axon projecting toward the medial or lateral olfactory tract [126]. Although the approaches employed for the analysis of *Drosophila* neurons and zebrafish mitral cells share significant similarities, we leveraged unique features of the mitral cells to inform the axon-dendrite segregation. Since the Mitral cells’ tufted dendritic structure is characterized by a high number of terminal points in comparison to the axonic region, we first define a rough dendritic compartment as the cluster yielded by the K-means method with the highest number of terminal points (**Figure S4B**, step 2). Refining the dendritic compartment in these cells presented additional challenges due to the occasional ambiguity in soma location from the reconstructions. To address this, we devised a new step in our method to identify the soma location. We identified the soma location as the node where all paths from the terminal nodes of the first-pass dendritic compartment, heading toward the axon terminal, converged. This allowed us to further segregate the dendritic compartment, providing a more nuanced and detailed understanding of the unique structure of zebrafish mitral cells (**Figure S4B**, step 3). Finally, to validate our approach, we confirmed our algorithm’s classifications through a consensus analysis conducted by two experts.

Finally, it is important to note that while our strategy offers a promising new approach for neuronal structure axon-dendrite annotation, there are a few limitations to consider. For example, the performance of the algorithm may be affected by the quality of the neuronal reconstructions and the choice of features used to represent each node. Furthermore, while we set K at 2 in this study, exploring different values of K could potentially provide more granular insights into neuronal architecture.

### Dendritic synaptic density calculations

In this study, we quantified dendritic synaptic densities as the ratio of the total number of synapses on the dendrite to its total dendritic cable length (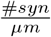; **Table 2**). We measured synaptic densities for all neurons in the species studied. For the *Drosophila* larva, and adult *Drosophila* and mouse datasets, the detection of synapses was virtually complete, eliminating the need for further proofreading or adjustments. For the zebrafish dataset, a correction of synaptic counts was necessary to address artifacts associated with electron microscopy (EM) reconstructions.

In particular, we developed an alternative approach to address artefacts in neuronal arbour reconstruction and synapse annotation, which impacted the measurement of dendritic synaptic density. During the reconstruction and proofreading of larval zebrafish neurons, certain biases were observed, which were attributed to the tracing and proofreading method employed in [45]. Specifically, arbour tracing began at the soma rather than at the synapses, leading to sparser reconstructions by an increased number of errors of omission (starting from the synapses would have provided more points of cable convergence onto a neuronal arbour). This bias was exacerbated by the application of the redundant skeleton consensus procedure (RESCOP; [127]). That is, multiple individuals reconstructed the same arbour, and then only the common subset of the traced cable was retained as a means to minimize reconstruction errors. Given that in dendrites, most errors are of omission and overwhelmingly impact thin calibre synapse-bearing terminal cable, the application of RESCOP in all likelihood removed a substantial number of its synapses from the resulting consensus arbours. Not only was the total number of synapses reduced, but also resulted in an over-representation of dendritic shafts, which are cables of known low synaptic density (**Figure S1A, B**). In order to address these biases and estimate the true dendritic synaptic density, we implemented a simple correction strategy for synaptic counts.The rationale is as follows: RESCOP-based reconstructions truncate terminal branches, which typically contribute minimal additional cable length but can contain 2–3 synapses [59]. To compensate this, we added 1, 2, or 3 synapses per terminal branch in each morphology to assess the effect on density estimates (**Figure S1C**). The recalculated synaptic densities under each correction level were as follows. Real data (no correction):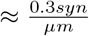; 1 added synapse per terminal: 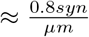; 2 added synapses: 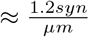; 3 added synapses: 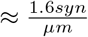. These corrected estimates span a biologically plausible range and show that even under the most aggressive correction, zebrafish synaptic density remains well within the interspecies range 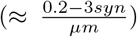. We selected the 1-synapse-per-terminal correction as our final adjustment, as a recent study reported that local synaptic density in zebrafish neurons falls between the levels predicted by the 1- and 2-synapse correction models [128]. We acknowledge that this is a rough estimate, but current evidence suggests it represents a lower-bound approximation of the true dendritic synaptic density (**Table 2**). Importantly, our sensitivity analysis demonstrates that these corrections do not affect the study’s central conclusion regarding the invariance of synaptic density across species.

For the adult *Drosophila* neurons, it was necessary to filter out data with low post-synaptic reconstruction completeness, as some neuronal arbours were only partially reconstructed. In particular, dendrites were often truncated (a false split), missing the small terminal synapse-bearing branches (the ‘twigs’ [59]), and therefore dendrites as provided present an over-representation of low-synapse density cable (the main shafts or ‘backbone’) in addition to potentially missing some synapses on the reconstructed cable. It is important to note that variability in postsynaptic reconstruction levels across brain regions is due to differences in the extent of proofreading, which led to varying estimates of missing synapses. Some regions, such as the central complex [41, 77] and mushroom body [129], were extensively proofread due to their relevance in respective studies, while others were not.

To estimate the impact of these reconstruction errors on dendritic synaptic density, we used the known amount of missing branches and synapses—referred to as the “postsynaptic completion rate”—which is accessible via neuprint.janelia.org, a tool for exploring connectomics datasets. neuprint.janelia.org. After splitting the morphologies into dendrites and axons, we calculated the ratio of the number of synapses to the total dendritic cable 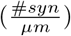 and compared it with the post-synaptic completion rate of the corresponding brain region. We found that the two quantities were correlated (*r* = 0.61, *p* < 0.001; **Figure S1D**). As a conservative measure to avoid biases in our analysis, we discarded all neurons in brain regions with a postsynaptic completion rate below 80%, using only neurons that satisfied this constraint in our analysis. For these neurons, no processing was performed. It is important to note that no brain region had a completion rate greater than 90%, which could potentially reduce the density values of the neurons analysed (**Table 3**).

### Dendritic synaptic density invariance and neuronal responses stabilisation

Neurons typically receive synaptic contacts across their dendritic tree. These dendrites, acting as leaky conductors, allow currents to propagate along dendritic cables towards the integration site [24]. At the same time, these afferent currents leak across the cell membrane. Intuitively, larger cells have lower input resistances due to their increased length and membrane surface area, which requires larger synaptic currents to achieve the same excitability levels as smaller neurons [23, 72]. However, larger dendrites can form more synaptic connections, boosting depolarisation due to increased afferent currents. In [29], it was theoretically demonstrated that these opposing phenomena precisely cancel each other out for any dendritic morphology, provided the dendritic synaptic density remains constant. Thus, a neuron’s voltage response (*V*_syn_) to distributed synaptic inputs are independent of total cable length and given by:

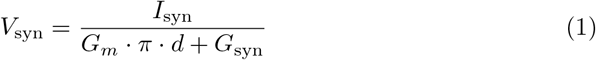

with *V*_syn_ the steady-state voltage response; *I*_syn_ is the total synaptic input current, representing the sum of currents from distributed synaptic inputs; *G*_*m*_ is the membrane conductance, reflecting the electrical properties of the dendritic membrane; *d* is the dendritic diameter; and *G*_syn_ is the total synaptic conductance.

This homeostatic effect ensures that, for a given synaptic density, neuronal voltage responses are determined solely by the average diameter of the dendritic and intrinsic conductivity [29]. The output voltage responses can then be approximated by *V*_*syn*_ ∝ *w*_*r*_, where *w*_*r*_ represents the relative connectivity of a given connection, defined as the percentage of activated dendritic synapses relative to the total number of dendritic postsynapses [29]. This relationship holds across a wide range of morphologies, regardless of their arborisation complexity, with the number of spikes closely approximating the relative weight of active synapses. This consistency occurs not only when synapses are uniformly activated but also when they are activated in clusters within a specific dendritic tree. Consequently, a neuron’s voltage response is largely dependent on synaptic density, highlighting the link between synaptic density invariance and functional stability.

### Dendritic inter-synaptic distance calculations

In our analysis, we computed the inter-synaptic distance by calculating the spatial locations of all synapses located along the paths extending from every terminal node leading to the dendritic root. Starting from the longest path to the root, we sequentially progressed through the longer paths until they reached the soma, or converged with a larger path length, at which point we transitioned to the next remaining longer path. The inter-synaptic distance was defined as the path length that stretches between one synapse and its immediate successor in the direction towards the dendritic root (**Table 2**). In the context of the zebrafish dataset, a simple correction was implemented. Reconstruction and proofreading biases inherent to this dataset can induce omission errors in synapse-dense regions, particularly within smaller terminal branches, potentially skewing the distance distribution towards larger values. Thus, the inter-synaptic distance for zebrafish was determined as described above, and then adjusted proportionally based on the corrected count of corrected synapses (**Figure S1**). This adjustment provides a closer estimation to the true inter-synaptic distance.

In general, this analysis was prompted by the hypothesis that although the density of the synaptic may be consistent between species, variations in the location of the synapse could significantly affect neuronal excitability. However, our analysis reveals that the inter-synaptic distance remains relatively constant across species, with *Drosophila* larva mean = 1.31*μm* ± 0.49; *Drosophila* adult mean = 1.89*μm* ± 0.51; zebrafish mean = 1.23*μm* ± 0.47 (uncorrected zebrafish inter-synaptic 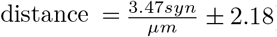); mouse mean = 1.57*μm* ± 0.45 (**Table 2**).

### Normalised Metric for Synapse Spatial Distribution

The spatial distribution of synapses on dendritic trees can influence the integration properties of neurons [23, 81, 90]. A non-uniform distribution, characterised by clusters of synapses located distally or proximally, can affect the way a neuron processes and integrates incoming signals. To analyse the synapse spatial distribution, we introduced a relative metric that evaluates the position of synapses regardless of dendritic morphology. It quantifies the relationship between the average path length location (PL) of the observed synapses and the expected PL distribution across a dendritic arbour, assuming a uniform random spatial distribution of synapses. In essence, the anatomical synapse locations are compared with the expected distribution of randomly sampled locations across all potential anatomical sites, specifically the nodes, within a given dendritic arbour. Here, the path length denotes the distance from a specific location in the dendritic tree to the root of the dendrite.

This metric provides a relative quantity, comparing the average observed synapse locations against the mean PL distribution of dendritic nodes after being sampled uniformly at random. In order to compare across different cell-types and species, we bound the measure between 0-1, based on the shortest (proximal - 0) and longest (distal - 1) PLs in a given dendritic tree, with 0.5 being the mean PL of dendritic nodes when sampled uniformly at random. Through this, we can determine whether the synapses are located primarily proximally or distally from the root compared to the typical spatial distribution of the dendritic nodes.

**Overall Synapse Distribution Metric**: the overall synapse distribution metric quantifies how the average PL of observed synapse location compares to the distribution of node PLs in the dendritic tree. The metric is computed as follows:

1. **Calculate the mean PL of observed synapses (*N*)**:

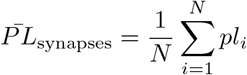
2. **Determine the bounds for normalization**: Let *D* represent the dataset of all dendritic nodes’ PL:

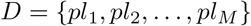

where *M* is the total number of nodes in the dendritic tree.
  - *Lower bound*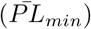: mean PL of the shortest *N* dendritic nodes in *D*.
  - *Upper bound* 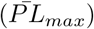: mean PL of the longest *N* dendritic nodes in *D*.
3. **Calculate the mean PL after random sampling**: Given *D*, we aim to compute the average PL from random samples. For each iteration *i* (with *i* = 1, 2, …, 10000), we randomly sample *N* nodes from *D* and compute the average PL for that sample:

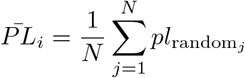

where *pl*_random_ is the *j*^*th*^ randomly sampled path length from *D* in the *i*^*th*^ iteration. Finally, after completing all 10000 iterations, we compute the overall average:

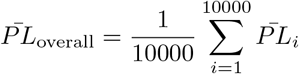

This 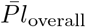 represents the average PL computed from 10000 random samples of *N* nodes each from the dataset *D*.
4. **Create a Mapping Function**: We used either linear interpolation or direct linear functions to compute normalised values based on given data points. Both methods establish a linear relationship between known points. The value we sought to map was the mean path length 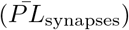 of observed synapses. By employing these techniques, we ensured that all values fall within the interval [0, 1]. Specifically, the randomly sampled mean serves as the breakpoint, mapping 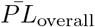 to 0.5. Additionally, the lower bound 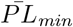 maps to 0, and the upper bound 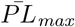 maps to 1. This approach results in a continuous and monotonic transformation, enabling a normalised comparison of synapse distribution within the neuron’s structure and facilitating comparison across different morphologies.

It is important to note that, while our methods provide a robust metric to analyse the spatial distribution of synapses in neurons, it is important to address specific dataset limitations. For the neurons of adult *Drosophila* included in our study, there exists a potential for under-sampling, as we have the smallest sample (*N* = 60). In the zebrafish dataset, reconstruction and proofreading biases can cause omission errors in synapse-rich areas, especially smaller terminal branches, potentially skewing our analysis towards a more proximal distribution. In fact, in our data, we observed that the spatial location of the mean synapse of the zebrafish is more skewed to the proximal locations. (*D*. larva mean = 0.55 ± 0.2; *D*. adult mean = 0.39 ± 0.12; zebrafish mean = 0.32 ± 0.15; mouse mean = 0.43 ± 0.2; **Table 2**).

### Analysis of synaptic density variance around the mean of 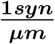

#### Synapse packing constraints

We wished to understand whether fluctuations around the mean of 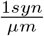 could be due to physical constraints [18, 128]. We reasoned that optimal packing tightly regulates the “volume budget” that synapses can occupy in a circuit. To test this, we used the MICrONS mm^3^ mouse dataset [47], the only available dataset in which synaptic sizes were quantified for all synapses (in arbitrary volume units derived from the voxel resolution of the EM volume) across all reconstructed cells (*N* = 1183; **Table 3**).

To assess how variation in synapse size influences estimated dendritic synaptic density, we implemented a strategy based on redistributing each neuron’s total synaptic volume using size-defined intervals from the empirical synapse size distribution. For each neuron *i*, we first computed the total synaptic volume:

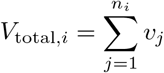

where *v*_*j*_ is the volume of the *j*-th synapse and *n*_*i*_ is the number of synapses on neuron *i*. We then used the global synapse size distribution (aggregated across all neurons) to define cumulative intervals—sorted either from the smallest to the largest synapses, or vice versa—in 5% increments. For each cumulative interval *k*, we computed the mean synapse size 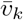. Assuming that the total synaptic volume remains fixed, the number of synapses that could be packed using the average size from interval *k* is:

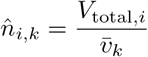

From this, we estimated the hypothetical synaptic density:

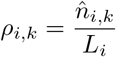

where *L*_*i*_ is the total dendritic length of neuron *i*. This approach allowed us to quantify a plausible range of synaptic densities that a neuron could exhibit under the same synaptic volume, but assuming different packing strategies biased toward smaller or larger synapses. These estimates formed upper and lower theoretical bounds, revealing how much density could vary purely due to synaptic size, independent of total synaptic volume or dendritic morphology (**Figure 4A, B**)

#### Correlation with synaptic size and dendritic radius

To correlate synaptic density with synaptic size and radii, we used mouse data sets (MICrONS mm^3^ and layer 2/3; **Table 1, 3**; [46, 47]). These were the only datasets that provided both synaptic size (*>* 97%; *N* = 1446) and radii measurements across most morphologies (100%; *N* = 1486), with data thoroughly proofread by expert annotators. Since synapse sizes were measured in arbitrary volume units in MICrONS mm^3^, and in real world units (*μm*^3^) in the layer 2/3 dataset, we normalised the synapse size data to provide a clearer interpretation of the results. Moreover, due to potential discrepancies introduced by different staining methods in mouse datasets, which could affect cable volume measurements due to shrinkage factors, we normalised the radii data for each mouse dataset [78].

### Testing wiring minimisation scaling law

To challenge the wire minimisation properties of dendritic trees in our dataset (*N* = 2486; **Table 3** for the datasets used), we verified if the total length of the dendrite (L), the number of dendrite synapses (n) and the volume (V) were in accordance with the scaling law, 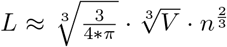, as proposed by [19] (**Figure 1A**). The aforementioned morphometrics were computed using built-in functions from the TREES Toolbox (MATLAB; https://www.treestoolbox.org). In particular, arbour volumes were computed using the `boundary_tree` function from the TREES Toolbox [83], following [130]. Here, which generates alpha shapes governed by a shrink factor (*S*), ranging from 0 (convex hull) to 1 (tightest connected boundary). For each neuron, the shrink factor was selected based on dendritic convexity following [130].

However, for the MICrONS mm^3^ dataset, convex and alpha shape analyses often overestimated volume in morphologies with sparse, colinear, or minimally branched geometries [130]. These structures leave large gaps between branches, causing boundary methods to wrap around empty space and distort the true spatial footprint [131]. To avoid these artifacts, we implemented a sparsity check. We classified neurons as highly sparse based on two biologically informed metrics: volumetric synaptic density in the convex hull 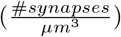 and length-to-volume (L/V) ratio 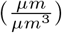. Specifically, we identified neurons with synaptic densities below 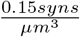 and L/V ratios smaller than 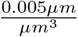 as outliers. These thresholds were motivated by comparisons with data reported in [132] and [19]. Neurons from the MICrONS mm^3^ dataset [47] consistently fell beyond both thresholds by more than an order of magnitude, prompting us to classify them as highly sparse. For these neurons, we applied a branchwise decomposition strategy, computing convex hulls separately for each dendritic branch to more accurately estimate their true occupied volume.

To ensure that our decomposition method did not trivially impose a length–volume scaling relationship (*L* ∼*V*), we performed two control simulations (**Figure S5**). First, we generated tortuous versions of the original MICrONS mm^3^ dataset morphologies. This was achieved by increasing the curvature of their branches without altering overall topology, using the `dscam_tree` function from the TREES toolbox. These modified trees showed a stronger correlation between length and volume (*r* = 0.535) than wild-type neurons (*r* = 0.32, *p <* 0.001). Second, we randomly reassigned terminal branches to new locations in dendritic morphology while preserving their individual geometry. This disrupted the overall spatial embedding and reduced the length–volume correlation (*r* = 0.187; *p <* 0.001) compared to wild type). Together, these results indicate that the observed length–volume scaling reflects genuine biological structure, not an artifact introduced by the decomposition method that forced *L* ∼ *V*.

To assess whether the empirically observed synaptic density of approximately 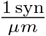 reflects pressure to optimise cable, we evaluated how varying synaptic density *ρ* affects the predictions from the scaling law. Because synapse number can be expressed as *n* = *ρ* · *L*, we substituted this into the equation to assess how deviations in assumed density *ρ* impact model predictions. For a given fixed *ρ*, we calculated predicted dendritic length *L*_pred_(*ρ*) for each neuron. To evaluate model accuracy, we computed the root-mean-square error (RMSE) between predicted and observed dendritic lengths:

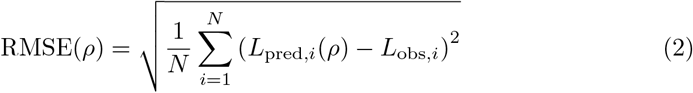

where *L*_obs,*i*_ is the observed dendritic length for neuron *i*, and *N* is the number of neurons. To make the error metric comparable across densities, we normalized the RMSE by the mean observed dendritic length:

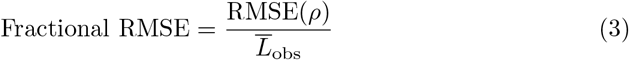

We systematically varied *ρ* by 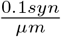 steps, over the observed range of 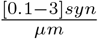. This range was motivated by the lower and upper limits of the synaptic densities observed in the real data (minimal observed 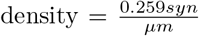, maximum observed 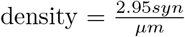).

### Biophysical passive steady-state models of cortical pyramidal cells

To investigate whether synaptic density invariance stabilises voltage responses across diverse dendritic morphologies, we used EM-constrained passive models based on morphologies from the MICrONS mm^3^ mouse dataset. This dataset provides high-resolution 3D reconstructions of cortical pyramidal neurons, ensuring the largest dataset with accurate dendritic diameter measurements. We did not include the MICrONS layer 2/3 dataset in the modeling. This decision was made to prevent confounding effects, as staining methods might influence the radius of morphologies across specimens, leading us to focus on the larger of the two available datasets [78].

Steady-state voltage responses (*V*_syn_) were calculated at the soma in response to activation of distributed synaptic inputs. Synaptic conductances in the models were scaled to reach a target voltage of 11 mV, selected to approximate the spiking threshold of mouse pyramidal cells [29]. Modelled morphologies were constrained with dendritic diameter and synaptic density from the MICrONS *mm*^3^ dataset, ensuring biologically representative parameters. Synapses were uniformly distributed across the dendritic arbour in each trial, justified by the observed spatial homogeneity of synaptic locations in the mouse data (**Table 2**). The number of synapses per neuron was determined from EM-derived synapse densities. The membrane conductance (*G*_*m*_ = 50*μ*S/cm^2^) was assumed constant, reflecting typical values for mouse pyramidal cells [29].

The total synaptic input current (*I*_syn_) was derived from the dendritic constancy equation, *V*_syn_ = *I*_syn_*/*(*G*_*m*_ ·*π* · *d* + *G*_syn_), initially omitting *G*_syn_ to estimate the input current needed to achieve a steady-state voltage of approximately 11 mV [29]. This simplification was necessary because *G*_syn_ depends on *I*_syn_, making the equation implicit and requiring an initial approximation. To account for the drop in input resistance introduced by synaptic conductance, we applied a small voltage overshoot (Δ*V*) during the initial current calculation. The overshoot was determined via the `compute_overshoot` function, which numerically identifies the smallest Δ*V* sufficient for a conductance-based input to reach the 11 mV target, given membrane properties and synaptic density. The total synaptic conductance was then calculated as *G*_syn_ = *I*_syn_*/E*_*e*_, assuming a reversal potential of *E*_*e*_ = 60 mV [29]. This value was distributed across all synapses. Finally, steady-state voltage responses were computed using the `syn_tree` function from the TREES toolbox (Matlab; [83]).

We evaluated three conditions using these models: (1) real dendritic diameters and cell-specific synaptic densities, (2) average diameters with real synaptic densities, and (3) real diameters with a fixed synaptic density of mouse 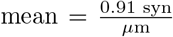. The grand mean dendritic diameter used for fixed-diameter simulations was diameter= 1.4*μ*m. Model responses were grouped into bins covering the full dendritic length range in 10% increments for comparison. These models were then compared against the dendritic constancy prediction, constrained with the same parameters as the models, with diameter and density set to the overall average.

### 3.6 Co-scaling of dendritic radius and PSD area

We correlated dendritic radii with synaptic size across species using four EM datasets: *Drosophila* larva [33], adult *Drosophila* [41, 77], and two mouse [46, 47]; and two LM datasets: rat [120] and human [119] (**Table 1**). To measure postsynaptic density areas, we used five EM datasets: *Drosophila* larva [33], adult *Drosophila* [122], mouse [71], and human [123] (**Table 4**).

To explore how dendritic radius and synaptic parameters interact to maintain stable voltage responses across species, we derived three models using the dendritic constancy equation. The simulated range of dendritic diameters (100–500*nm*) reflects the full observed range across species, with the initial value (100*nm*) chosen to represent the smallest brain in our dataset, similar to the *Drosophila* larva. Voltage responses in all cases were normalised to this initial condition.

In the first condition, synaptic input current (*I*_syn_) and synaptic conductance (*G*_syn_) scaled linearly with dendritic diameter (*d*), based on the observed relationship between radius and synaptic density across species (**Figure 6B**). The membrane conductance *G*_*m*_ was assumed constant for simplicity. The voltage response was calculated as:

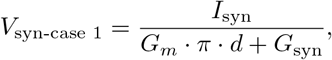

where *I*_syn_ and *G*_syn_ were derived using a the linear relationship between radius and PSD area across species (**Figure 6B**). This case represents a scenario where dendritic size scaling is accompanied by proportional increases in synaptic inputs and conductance. Voltage responses were normalised to the initial response at *d* = 100 nm.

In the second case, the dendritic diameter was held constant at its minimum value (*d*_min_ = 100 nm), while *I*_syn_ and *G*_syn_ scaled linearly with dendritic diameter, as in case 1. The voltage response was computed as:

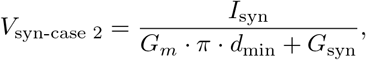

reflecting a situation where synaptic parameters grow with dendritic size, but the dendrite’s radius remains constrained. Voltage responses were normalised by the initial response.

Finally, in the third case dendritic diameter (*d*) scaled across the range, while *I*_syn_ and *G*_syn_ were fixed at their initial values (*I*_syn-fixed_ = *I*_syn_, for *d* = 100. The voltage response was calculated as:

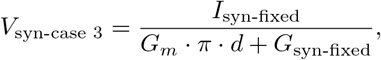

which simulates dendritic growth without corresponding increases in synaptic inputs. Normalisation was applied to the initial condition.

### Statistical analysis

Statistical analyses were performed using Navis (Python), Natverve (R), and TREES Toolbox (Matlab). We used Pearson’s correlation and Spearman rank correlation to analyse the relationships between various parameters in our datasets. Permutation tests were used to quantify significance levels (10000 permutations). The number of observations, significance values, and other relevant information for data comparisons are specified in the respective figure legend and in text. In all figures, * represents p value *<* 0.05, ** represents p value *<* 0.01, and *** represents p value *<* 0.001.

## Supplementary figures

**Fig. S1.**
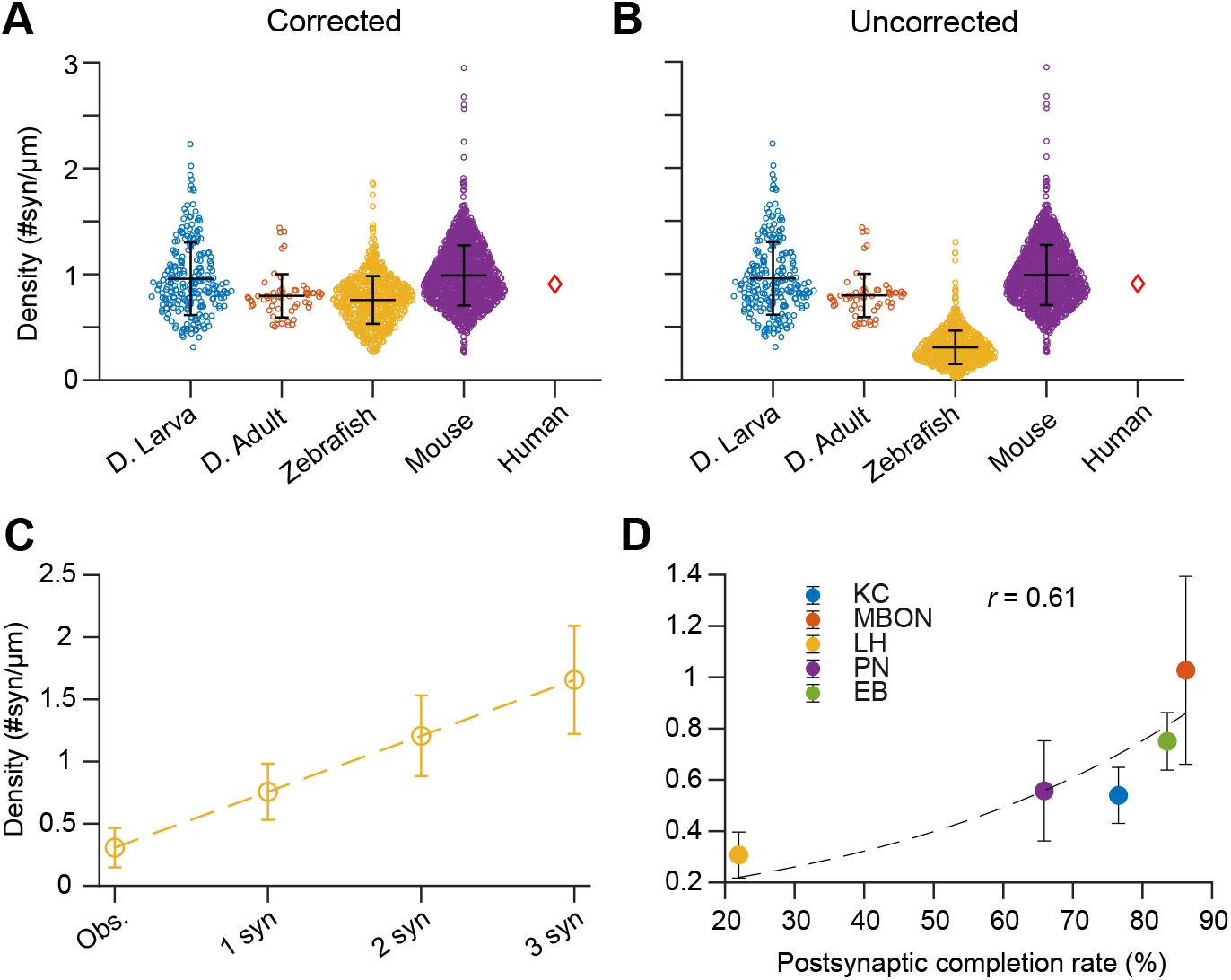
Synaptic density corrections and correlation analysis with postsynaptic reconstruction completeness level. **A**, Same as **Figure 1C. B**, Similar swarm plot of synaptic densities across species as in **A**, but with uncorrected zebrafish density values (see Methods 3.5). **C** Zebrafish synaptic density estimates following terminal-branch corrections. Observed synaptic densities (Obs.) were recalculated after adding 1, 2, or 3 synapses per terminal branch. Each point represents the mean synaptic density 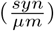 across zebrafish neurons, and error bars indicate the standard deviation. **D**, Correlation plot showing dendritic synaptic density as a function of postsynaptic completion rate in the adult fly Hemibrain connectome (*ρ* = 0.61, *p <* 0.001; Table 1, 2; see 3.5) [41, 77]. Different colours indicate different cell types in the adult fly brain. Error bars indicate the standard deviation. The data are fitted with an exponential fit (black dashed line) to assess the relationship between reconstruction completion rate and synaptic density. The correlation coefficient (*r*) was obtained by Spearman’s rank correlation, and *p* values were obtained from a permutation test (10000 permutations).

**Fig. S2.**
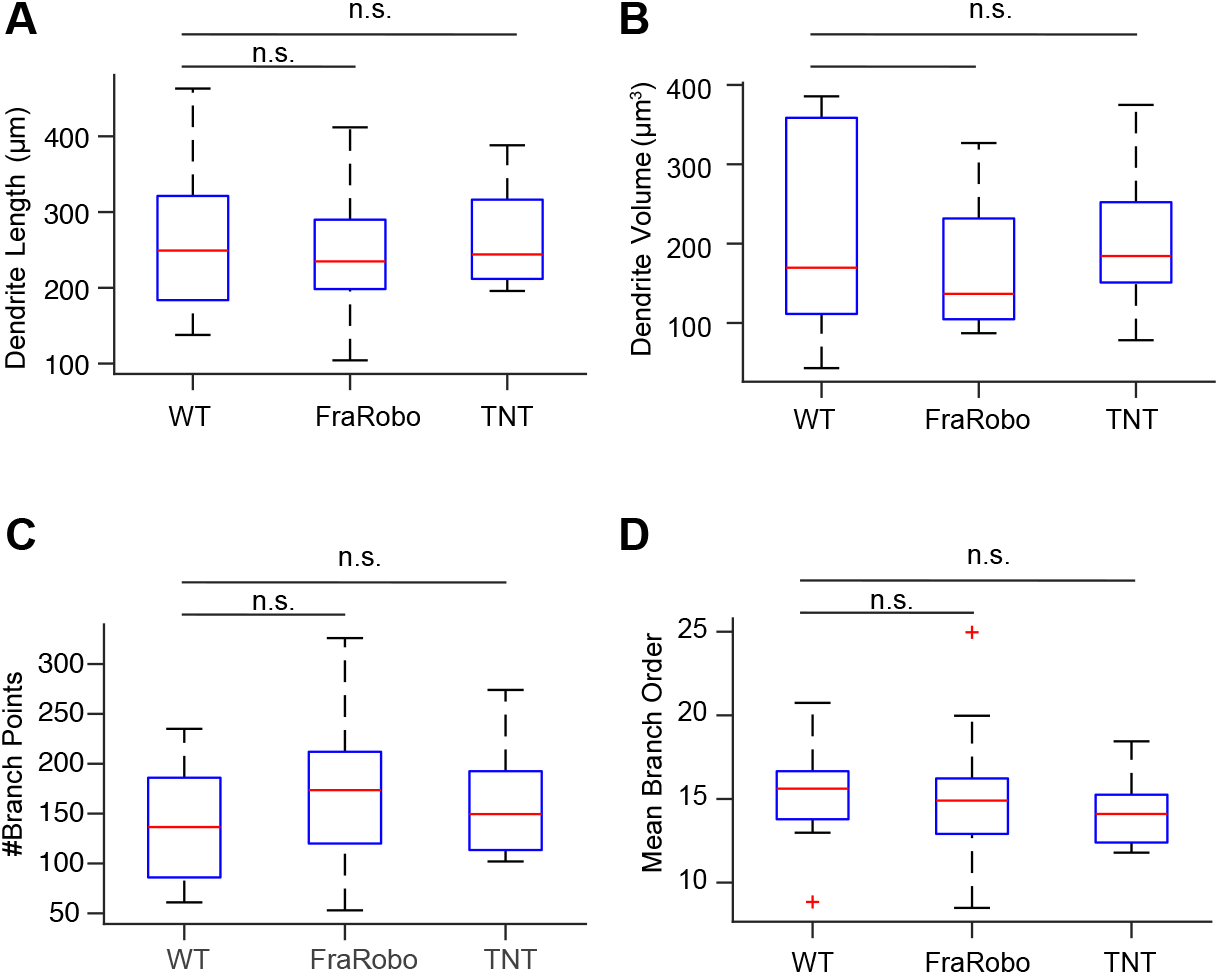
Comparative analysis of dendritic morphometrics in WT, FraRobo, and TNT EM volumes. **A**, Box plot showing the distribution of dendritic length across wild type (WT; *N* = 14), FraRobo (*N* = 14), and TNT (*N* = 12) mutants. **B-D**, same as **A**, but for dendritic volume, number of branch points, and mean branch order measurements. The red plus symbols indicate outliers within the data set for each condition. Outliers were defined based on the interquartile range. Horizontal lines above the plots denote statistical comparisons, with “n.s.” indicating a lack of statistical significance (*p >* 0.05), with *p* values obtained from permutation tests (10000 permutations).

**Fig. S3.**
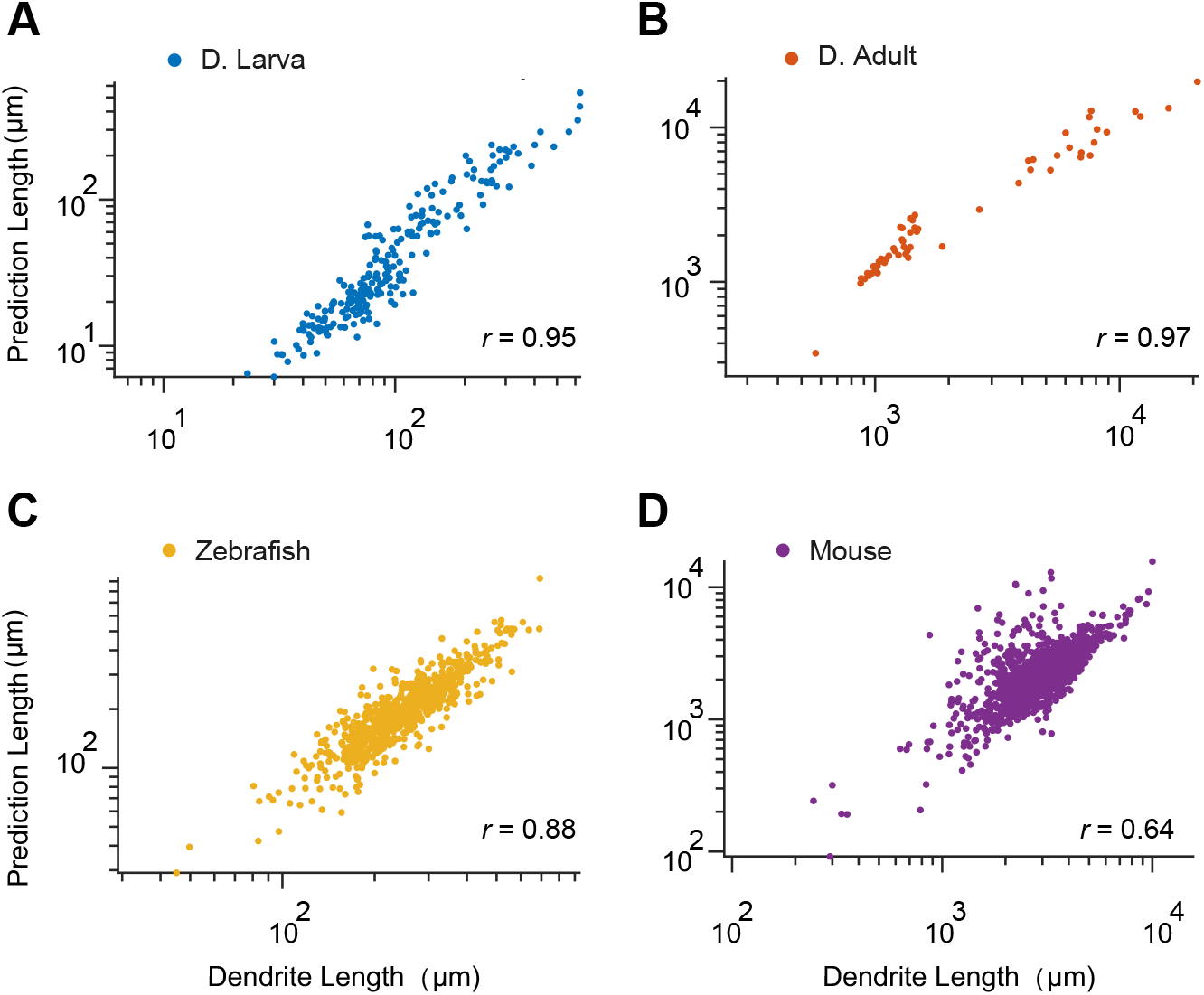
Scaling relationships plotted across individual species. The data is the same as in Figure 3C. **A**, Shown is the relation between total dendrite length of real dendrites (*L*_*Real*_) and the predicted length (*L*_*Predicted*_) computed using the scaling law for all *Drosophila* larva morphologies (*N* = 233; *r* = 0.95, *p <* 0.001; log-transformed data; Table 1). **B-D**, same as in **A** but for adult *Drosophila* (*N* = 60; *r* = 0.94, *p <* 0.001); larval zebrafish (*N* = 709; *r* = 0.9, *p <* 0.001); and mouse (*N* = 1484; *r* = 0.64, *p <* 0.001). Correlation coefficients (*r*) were obtained by Pearson correlation, and *p* values were obtained from a permutation tests (10000 permutations).

**Fig. S4.**
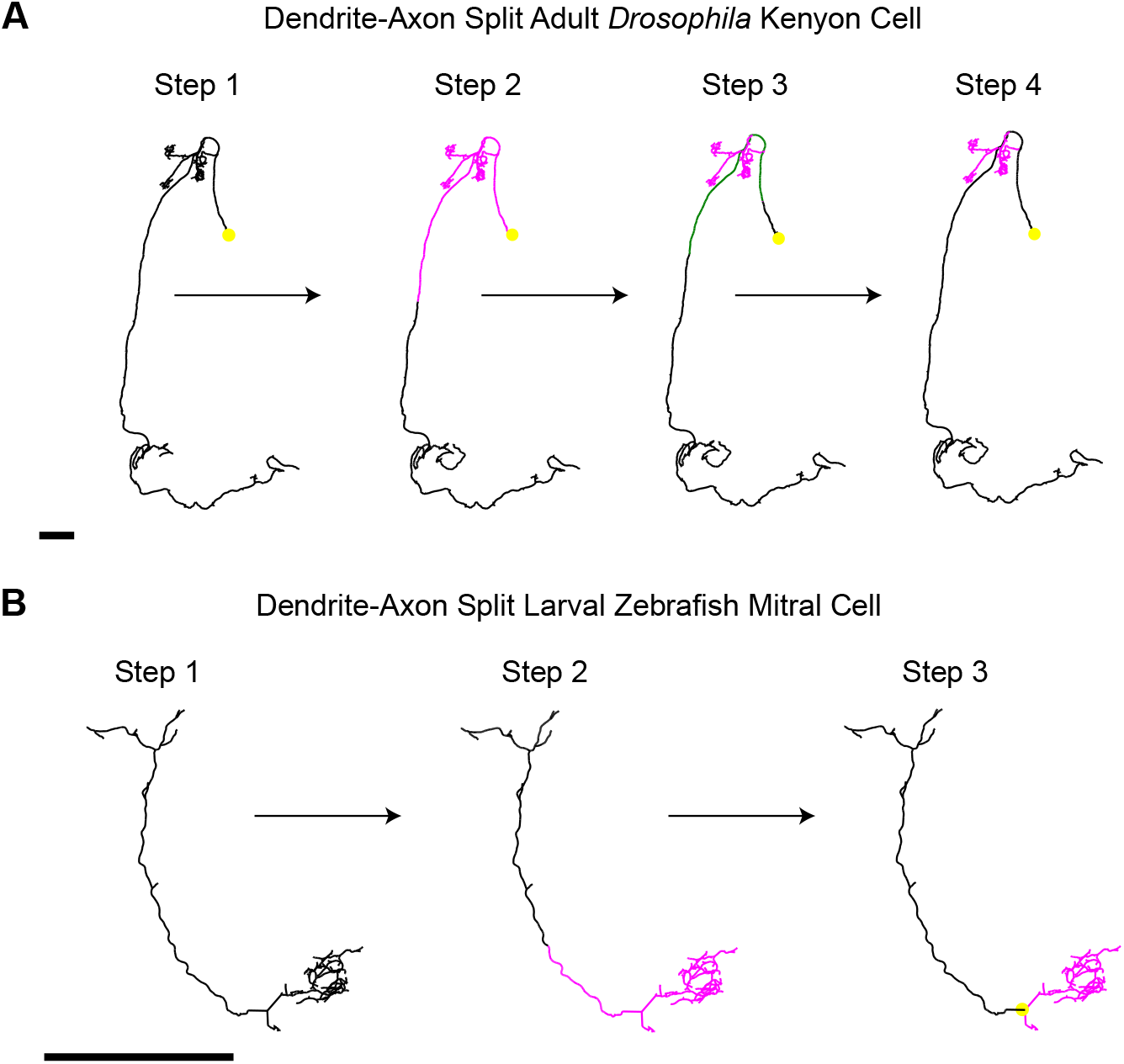
Illustrative dendrite-axon split algorithm for adult *Drosophila* and zebrafish datasets. **A**, Dendrite-axon split in adult *Drosophila* Kenyon-cell. The figure shows the four-step process: starting with the reconstructed morphology, then illustrating the separated compartments using K-means, highlighting the dendritic (pink) and axonal (black) and main trunk (green), and finally presenting the dendritic tree after main branch removal, with the soma marked (yellow dot). **B**, Dendrite-axon split of larval zebrafish mitral cell. The three-step process starts with representative morphology, then shows separated compartments (dendritic in pink and axonal in black) using K-means, and finally highlights the dendritic compartment (pink) and the soma proxy location (yellow). Scale bars: 10 *μm*.

**Fig. S5.**
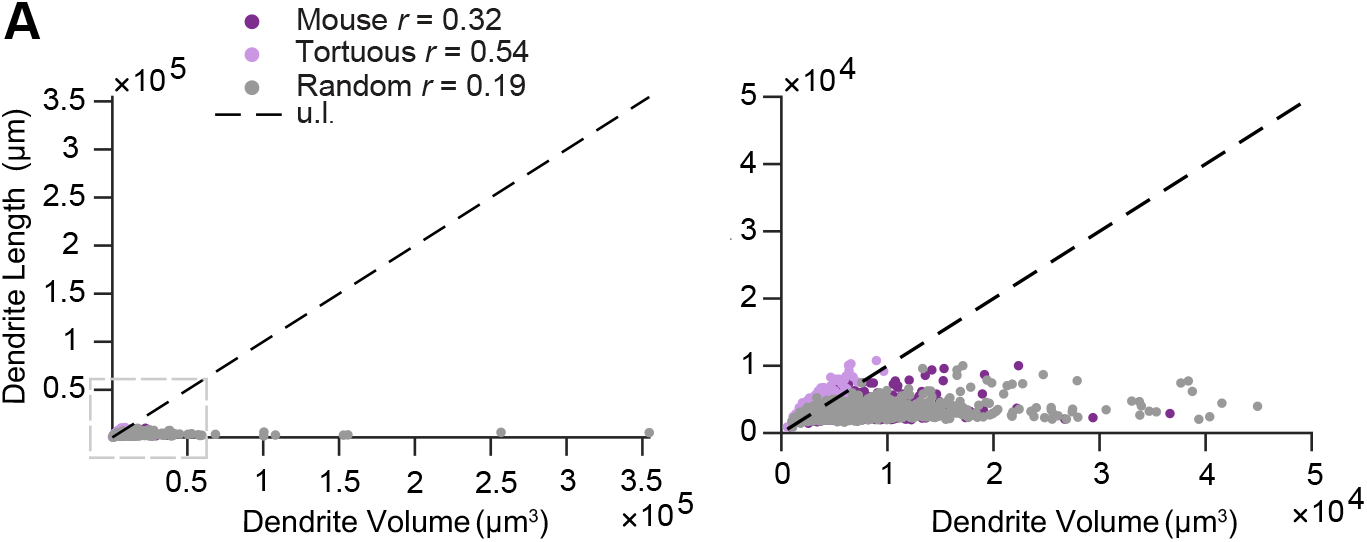
Simulations show that observed dendrite length–volume scaling is not imposed by decomposition method in MICrONS mm^3^ dataset. **A**, Relationship between dendrite length and dendrite volume for three groups (*N* = 1183): wild-type mouse neurons (purple; MICrONS mm^3^ dataset), tortuous morphologies (light purple), and randomly reconnected morphologies (dark gray). The dashed line indicates the unity line (*slope* = 1). (Right) Zoom-in of the region within the gray rectangle to highlight differences among groups. All correlations are statistically significant (*p <* 0.001). Correlation coefficients (*r*) were obtained using Pearson correlation, and *p* values obtained from a permutation test (10000 permutations)

